# Estimating temporally variable selection intensity from ancient DNA data

**DOI:** 10.1101/2022.08.01.502345

**Authors:** Zhangyi He, Xiaoyang Dai, Wenyang Lyu, Mark Beaumont, Feng Yu

## Abstract

Novel technologies for recovering DNA information from archaeological and historical specimens have made available an ever-increasing amount of temporally spaced genetic samples from natural populations. These genetic time series permit the direct assessment of patterns of temporal changes in allele frequencies, and hold the promise of improving power for the inference of selection. Increased time resolution can further facilitate testing hypotheses regarding the drivers of past selection events such as the incidence of plant and animal domestication. However, studying past selection processes through ancient DNA (aDNA) still involves considerable obstacles such as postmortem damage, high fragmentation, low coverage and small samples. To circumvent these challenges, we introduce a novel Bayesian framework for the inference of temporally variable selection based on genotype likelihoods instead of allele frequencies, thereby enabling us to model sample uncertainties resulting from the damage and fragmentation of aDNA molecules. Also, our approach permits the reconstruction of the underlying allele frequency trajectories of the population through time, which allows for a better understanding of the drivers of selection. We evaluate its performance through extensive simulations and demonstrate its utility with an application to the ancient horse samples genotyped at the loci for coat colouration. Our results reveal that incorporating sample uncertainties can further improve the inference of selection.

## 1. Introduction

A problem of long standing in population genetics is to understand evolutionary processes, including selection, in shaping the genetic composition of populations. Although the majority of these studies have focused on single time point polymorphism data, the use of temporally spaced genetic samples can provide valuable information on selection and demography (Bank et al., 2014; Skoglund et al., 2014). The commonest data sources have been evolve and resequencing studies combining experimental evolution under controlled laboratory or field mesocosm conditions with next-generation sequencing technology, which however are typically limited to the species with small evolutionary timescales (*e.g*., Turner & Miller, 2012; Bosshard et al., 2017; Good et al., 2017). Recent advances in technologies for obtaining DNA molecules from ancient biological material have resulted in massive increases in time serial samples of segregating alleles from natural populations (*e.g*., Mathieson et al., 2015; Librado et al., 2017; Fages et al., 2019), which offer unprecedented opportunities to study the chronology and tempo of selection across evolutionary timescales (see Dehasque et al., 2020, for a review).

As the number of published ancient genomes is growing rapidly, a range of statistical methods that estimate selection coefficients and other population genetic parameters based on ancient DNA (aDNA) data have been introduced over the last fifteen years (*e.g*., Bollback et al., 2008; Malaspinas et al., 2012; Mathieson & McVean, 2013; Steinrücken et al., 2014; Foll et al., 2014, 2015; Shim et al., 2016; Schraiber et al., 2016; Ferrer-Admetlla et al., 2016; He et al., 2020a,b; Mathieson, 2020; Lyu et al., 2022). See Malaspinas (2016) for a review. Most existing methods are built upon the hidden Markov model (HMM) framework of Williamson & Slatkin (1999), where the latent population allele frequency is modelled as a hidden state following the Wright-Fisher model (Fisher, 1922; Wright, 1931), and at each sampling time point, the observed sample allele frequency drawn from the underlying population is modelled as a noisy observation of the latent population allele frequency. To remain computationally feasible, these approaches usually work with the diffusion approximation of the Wright-Fisher model (*e.g*., Bollback et al., 2008; Malaspinas et al., 2012; Steinrücken et al., 2014; Schraiber et al., 2016; Ferrer-Admetlla et al., 2016; He et al., 2020a,b; Lyu et al., 2022). The diffusion approximation enables us to efficiently integrate over all possible underlying allele frequency trajectories, therefore producing substantial reductions in the computational cost of likelihood calculation. These methods have already been successfully applied in aDNA studies (*e.g*., Ludwig et al., 2009; Sandoval-Castellanos et al., 2017; Ye et al., 2017; Wutke et al., 2018). Moment-based approximations of the Wright-Fisher model, as tractable alternatives, are commonly used in the approaches tailored to experimental evolution (*e.g*., Lacerda & Seoighe, 2014; Feder et al., 2014; Terhorst et al., 2015; Paris et al., 2019) due to their poor performance for large evolutionary timescales (He et al., 2021).

Although the field of aDNA is experiencing an exponential increase in terms of the amount of available data, accompanied by an increase in available tailored statistical approaches, data quality remains a challenge due to postmortem damage, high fragmentation, low coverage and small samples. To our knowledge, none of the existing methods allow for modelling the two main characteristics of aDNA, *i.e*., the high error rate caused by the damage of aDNA molecules and the high missing rate resulting from the fragmentation of aDNA molecules, with the exception of Ferrer-Admetlla et al. (2016) and He et al. (2020a), which partially resolved this issue. Ferrer-Admetlla et al. (2016) incorporated genotype calling errors but excluded samples with missing genotypes while He et al. (2020a) allowed genotype missing calls but assumed no calling errors. Moreover, most existing approaches assume that the selection coefficient is fixed over time, which is commonly violated in aDNA studies. Most cases of adaptation in natural populations involve adaptation to ecological, environmental and cultural shifts, where it is no longer appropriate for the selection coefficient to remain constant through time. Shim et al. (2016) extended WFABC (Foll et al., 2015) to detect and quantify changing selection coefficients from genetic time series but more suitable for the scenario of a short evolutionary timescale like experimental evolution due to computational efficiency. More recently, Mathieson (2020) introduced a novel method to infer selection and its strength and timing of changes from aDNA data. However, these methods still ignored genotype calling errors and missing calls.

To address these challenges, in this work we introduce a novel Bayesian approach for estimating temporally variable selection intensity from aDNA data while modelling sample uncertainties arising from postmortem damage, high fragmentation, low coverage and small samples. We test our method through extensive simulations, in particular when samples are sparsely distributed in time with small sizes and in poor quality (*i.e*., high missing rate and error rate). We reanalyse a dataset of 201 ancient horse samples from Wutke et al. (2016) that were genotyped at the loci encoding coat colouration to illustrate the applicability of our approach on aDNA data.

## 2. New Approaches

To model sample uncertainties caused by the damage and fragmentation of aDNA molecules, unlike existing approaches, we base our selection inference on raw reads rather than called genotypes. Our method is built upon a novel two-layer HMM framework, where the first hidden layer characterises the underlying frequency trajectory of the mutant allele in the population through the Wright-Fisher diffusion, the second hidden layer represents the unobserved genotype of the individual in the sample, and the third observed layer represents the data on aDNA sequences (see Figure 1 for the graphical representation of our two-layer HMM framework). Such an HMM framework enables us to work with genotype likelihoods as input rather than allele frequencies. Our posterior computation is carried out by applying the particle marginal Metropolis-Hastings (PMMH) algorithm introduced by Andrieu et al. (2010), which permits a joint update of the selection coefficient and population mutant allele frequency trajectory. The reconstruction of the underlying population allele frequency trajectories allows for a better understanding of the drivers of selection. Moreover, our approach provides a procedure for testing hypotheses about whether the shift in selection is linked to specific ecological, environmental and cultural drivers. Further details on our method appear in Section 5.

**Figure 1:**
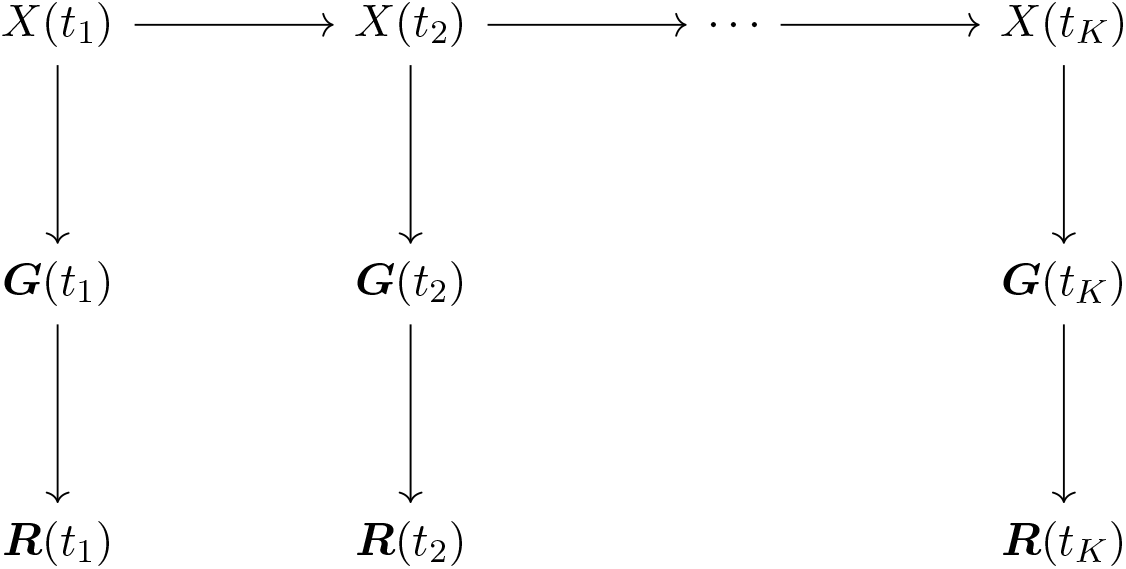
Graphical representation of our two-layer HMM framework for the data on aDNA sequences, where *X* denotes the population mutant allele frequency, ***G*** denotes the sample individual genotypes, and ***R*** denotes the sample individual reads.

## 3. Results

In this section, we test our approach through extensive simulations and show its applicability with the data on aDNA sequences from earlier studies of Ludwig et al. (2009), Pruvost et al. (2011) and Wutke et al. (2016), where they sequenced a total of 201 ancient horse samples for eight loci that determine horse coat colouration. In what follows, we base our selection inference on the maximum *a posteriori* probability (MAP) estimate and assume only a single event that might change selection.

### 3.1. Performance evaluation

We run forward-in-time simulations of the Wright-Fisher model with selection (*e.g*., Durrett, 2008) and assess the performance of our method with simulated datasets of genotype likelihoods. We let the only event that might change selection occur in generation 350, thereby taking the selection coefficient to be *s*(*k*) = *s*^−^ for *k <* 350 otherwise *s*(*k*) = *s*^+^. We uniformly draw the selection coefficients *s*^−^ and *s*^+^ from [−0.05, 0.05] and pick the dominance parameter of *h* = 0.5 (*i.e*., assuming codominance). To mimic the demographic history of the horse population given in Der Sarkissian et al. (2015), we adopt a bottleneck demographic history, where the population size *N* (*k*) = 32000 for *k <* 200, *N* (*k*) = 8000 for 200 ≤ *k <* 400, and *N* (*k*) = 16000 for *k* ≥ 400. The initial population mutant allele frequency *x*_1_ is uniformly drawn from [0.1, 0.9]. For clarity, we write down the procedure of generating the dataset of genotype likelihoods:

Repeat Step 1 until *x*_*K*_ ∈ (0, 1):

Step 1: Generate *s*^−^, *s*^+^ and ***x***_1:*K*_.

Step 1a: Draw *s*^−^, *s*^+^ from a uniform distribution over [−0.05, 0.05] and *x*_1_ from a uniform distribution over [0.1, 0.9].

Step 1b: Simulate ***x***_1:*K*_ with *s*^−^, *s*^+^ and *x*_1_ through the Wright-Fisher model with selection. Repeat Step 2 for *k* = 1, 2, …, *K*:

Step 2: Generate *p*(***r***_*n,k*_ | *g*) for *g* = 0, 1, 2 and *n* = 1, 2, …, *N*_*k*_:

Step 2a: Draw *g*_*n,k*_ with probabilities

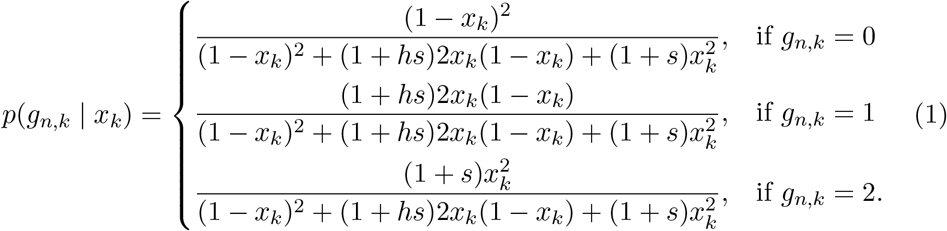

Step 2b: Draw *p*(***r***_*n,k*_ | *g*) for *g* = 0, 1, 2 from a Dirichlet distribution of order 3 with 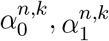 and 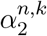.

We take the parameter 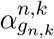 to be *ϕψ* and the other two to be (1−*ϕ*)*ψ/*2, where *ϕ* and *ψ* are the parameters introduced to control the quality of the simulated dataset in terms of the missing rate and error rate with a common threshold for genotype calling (*i.e*., 10 times more likely, see Kim et al., 2011). By following this procedure, we simulate a total of 801 generations starting from generation 0 and draw a sample of 10 individuals every 40 generations, 210 sampled individuals in total (*i.e*., nearly the size of the ancient horse samples), in our simulation studies.

We run a group of simulations to assess how our method performs for different data qualities, where we vary the parameters *ϕ* ∈ {0.75, 0.85, 0.95} and *ψ* ∈ {0.5, 1.0}, respectively, giving rise to six possible combinations of the missing rate and error rate (see Table 1 where the mean and standard deviation of missing rates and error rates are computed with 1000 replicates for each combination). For each scenario listed in Table 1, we consider nine possible combinations of the selection coefficients *s*^−^ and *s*^+^ (see Table 2), and for each combination we repeatedly run the procedure described above until we get 200 datasets of genotype likelihoods. Thus, in summary, we consider 200 replicates for each of 54 combinations of the data quality and selection scenario.

**Table 1:** Summary of data qualities across different combinations of the parameters *ϕ* and *ψ*.

| Scenario | $\phi$ | $\psi$ | Missing rate | | Error rate | |
| --- | --- | --- | --- | --- | --- | --- |
|  |  |  | Mean | Std. Dev. | Mean | Std. Dev. |
| A | 0.75 | 0.50 | 0.3095 | 0.0323 | 0.1392 | 0.0278 |
| B | 0.75 | 1.00 | 0.4542 | 0.0357 | 0.0711 | 0.0229 |
| C | 0.85 | 0.50 | 0.2037 | 0.0292 | 0.0673 | 0.0192 |
| D | 0.85 | 1.00 | 0.3029 | 0.0319 | 0.0275 | 0.0140 |
| E | 0.95 | 0.50 | 0.0753 | 0.0182 | 0.0172 | 0.0094 |
| F | 0.95 | 1.00 | 0.1108 | 0.0216 | 0.0059 | 0.0057 |

**Table 2:**
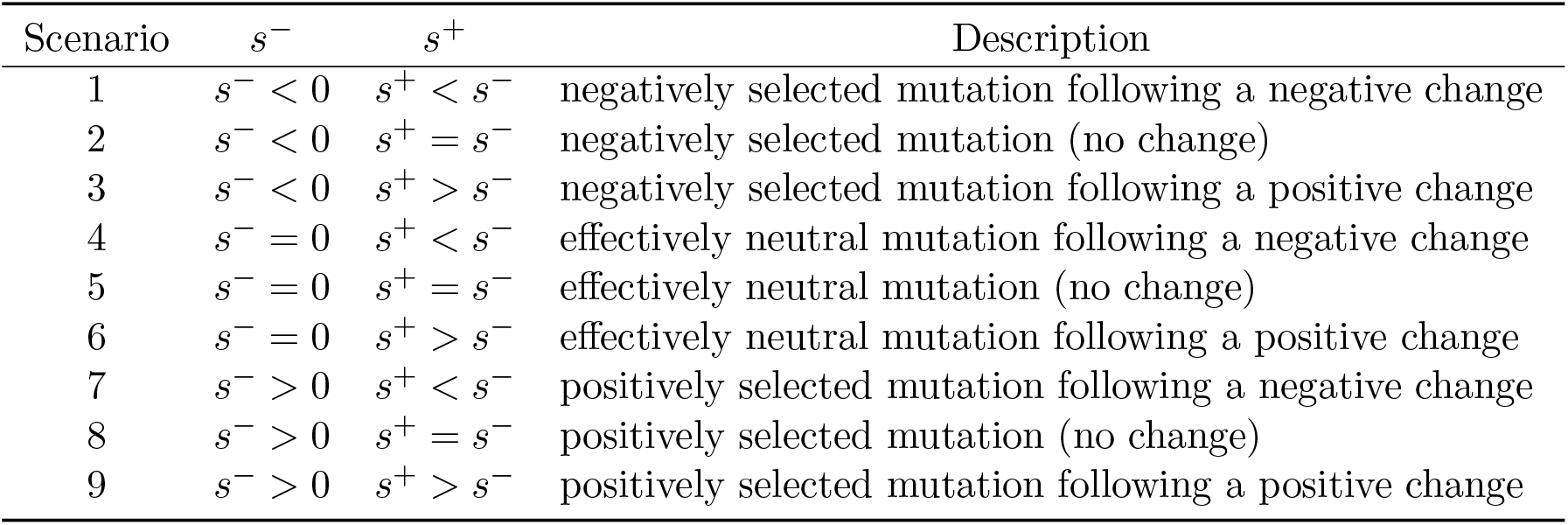
Summary of selection scenarios across different combinations of the selection coefficients *s*^−^ and *s*^+^.

| Scenario | $s^-$ | $s^+$ | Description |
| --- | --- | --- | --- |
| 1 | $s^- < 0$ | $s^+ < s^-$ | negatively selected mutation following a negative change |
| 2 | $s^- < 0$ | $s^+ = s^-$ | negatively selected mutation (no change) |
| 3 | $s^- < 0$ | $s^+ > s^-$ | negatively selected mutation following a positive change |
| 4 | $s^- = 0$ | $s^+ < s^-$ | effectively neutral mutation following a negative change |
| 5 | $s^- = 0$ | $s^+ = s^-$ | effectively neutral mutation (no change) |
| 6 | $s^- = 0$ | $s^+ > s^-$ | effectively neutral mutation following a positive change |
| 7 | $s^- > 0$ | $s^+ < s^-$ | positively selected mutation following a negative change |
| 8 | $s^- > 0$ | $s^+ = s^-$ | positively selected mutation (no change) |
| 9 | $s^- > 0$ | $s^+ > s^-$ | positively selected mutation following a positive change |

For each replicate, we choose a uniform prior over [−1, 1] for both selection coefficients *s*^−^ and *s*^+^, and adopt the reference population size *N*_0_ = 16000. We run 10000 PMMH iterations with 1000 particles, where each generation is partitioned into five subintervals in the Euler-Maruyama scheme. We discard the first half of the total PMMH samples as burn-in and thin the remaining by keeping every fifth value. See Figure 2 for our posteriors for the selection coefficients *s*^−^ and *s*^+^ produced from a simulated dataset of genotype likelihoods (see Supplementary Material, Table S1), including our estimate for the underlying frequency trajectory of the mutant allele in the population. Evidently, in this example our approach can accurately infer temporally variable selection from genetic time series in genotype likelihood format. The true underlying frequency trajectory of the mutant allele in the population fluctuates slightly around our estimate and is completely covered in our 95% highest posterior density (HPD) intervals.

**Figure 2:**
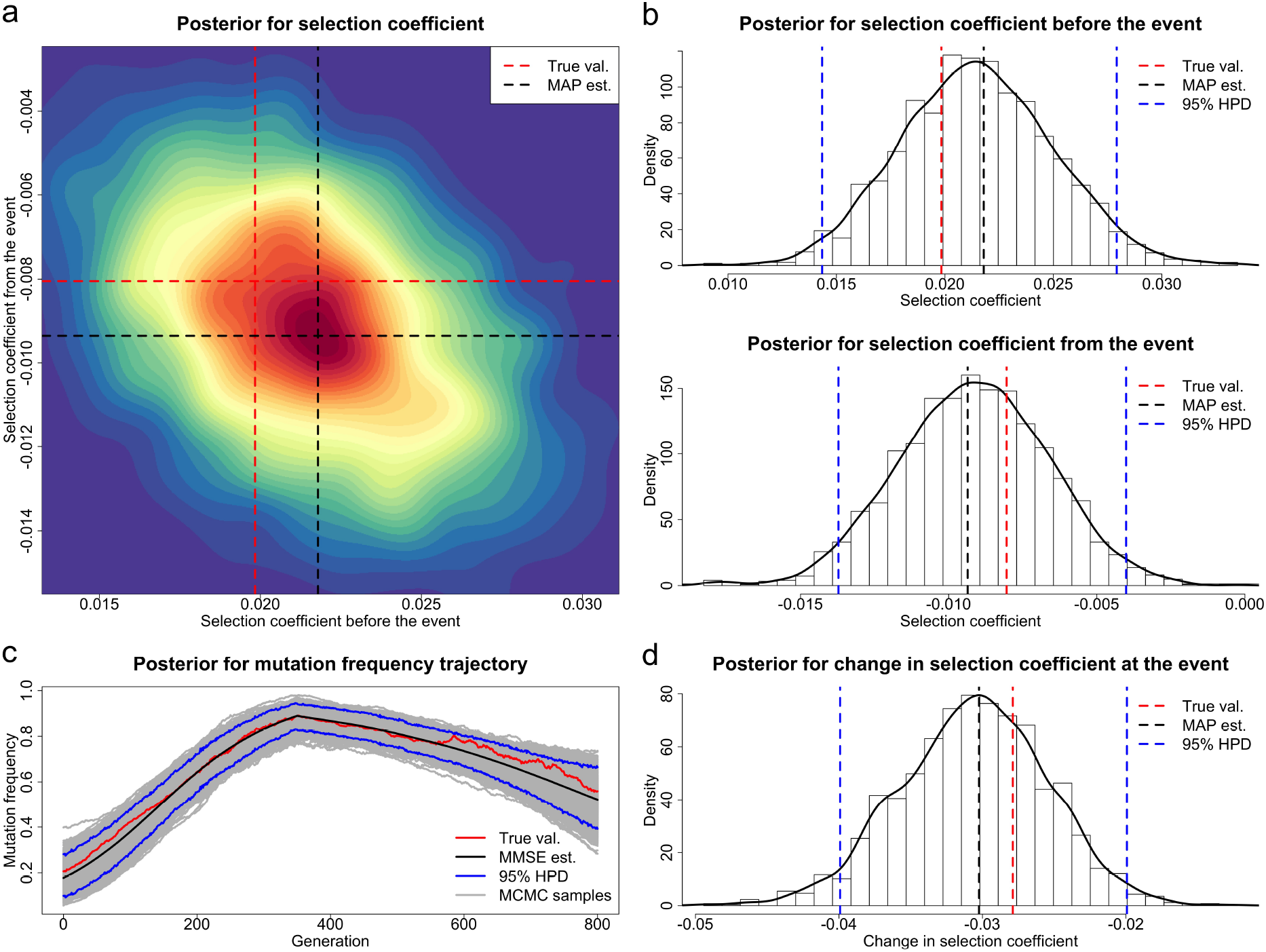
Posteriors for the selection coefficients and the underlying frequency trajectory of the mutant allele in the population produced through our method from a dataset of genotype likelihoods generated with the selection coefficients *s*^−^ = 0.0198 and *s*^+^ = −0.0081. (a) Joint posterior for the selection coefficients *s*^−^ and *s*^+^. (b) Marginal posteriors for the selection coefficients *s*^−^ and *s*^+^. (c) Posterior for the underlying trajectory of the mutant allele frequency. (d) Posterior for the selection change Δ*s*.

### 3.1.1. Performance in estimating selection coefficients

To assess how our method performs for estimating selection coefficients, we show the boxplot results of our estimates across different data qualities in Figure 3, where the tips of the whiskers are the 2.5%-quantile and the 97.5%-quantile, and the boxes denote the first and third quartiles with the median in the middle. We summarise the bias and root mean square error (RMSE) of our estimates in Supplementary Material, Table S2. As illustrated in Figure 3, our estimates for both selection coefficients are nearly median-unbiased across different data qualities although a slightly large bias can be found when the data quality is poor such as scenario A (around 31.0% missing rate and 13.9% error rate). The bias completely vanishes as the data quality improves (see, *e.g*., scenario F).

**Figure 3:**
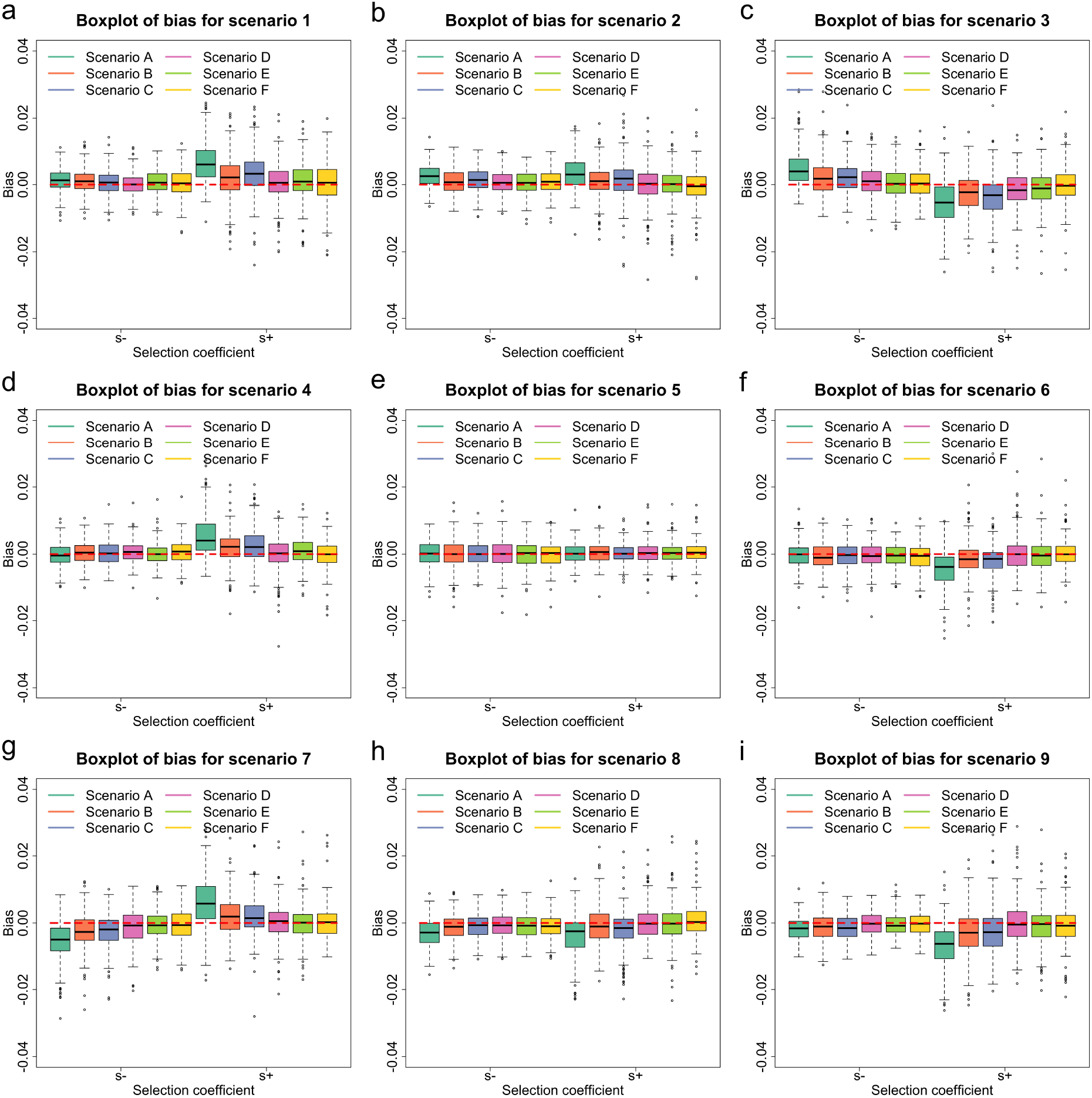
Empirical distributions for the bias in MAP estimates of the selection coefficients across different data qualities and selection scenarios. Data qualities (scenarios A–F) are described in Table 1, and selection scenarios (scenarios 1–9) are described in Table 2. (a)–(i) Boxplots of the bias for scenarios 1–9.

To assess how our method performs for different levels of selection coefficient, especially weak selection, we run an additional group of simulations with an example of codominance, where we adopt the parameters *ϕ* = 0.85 and *ψ* = 1. We assume no event that changes selection and vary the selection coefficient *s* ∈ [−0.05, 0.05], which is divided into nine subintervals [−0.05, −0.01), [−0.01, −0.005), [−0.005, −0.001), [−0.001, 0), {0}, (0, 0.001], (0.001, 0.005], (0.005, 0.01] and (0.01, 0.05]. For each subinterval, *e.g*., [−0.01, −0.005), we uniformly choose the selection coefficient *s* from [−0.01, −0.005), with which we generate a dataset of genotype likelihoods. Repeat this procedure until we obtain 200 simulated datasets for each subinterval. We show the boxplot results of our estimates across different levels of selection coefficient in Figure 4 and summarise the bias and RMSE of our estimates in Supplementary Material, Table S3.

**Figure 4:**
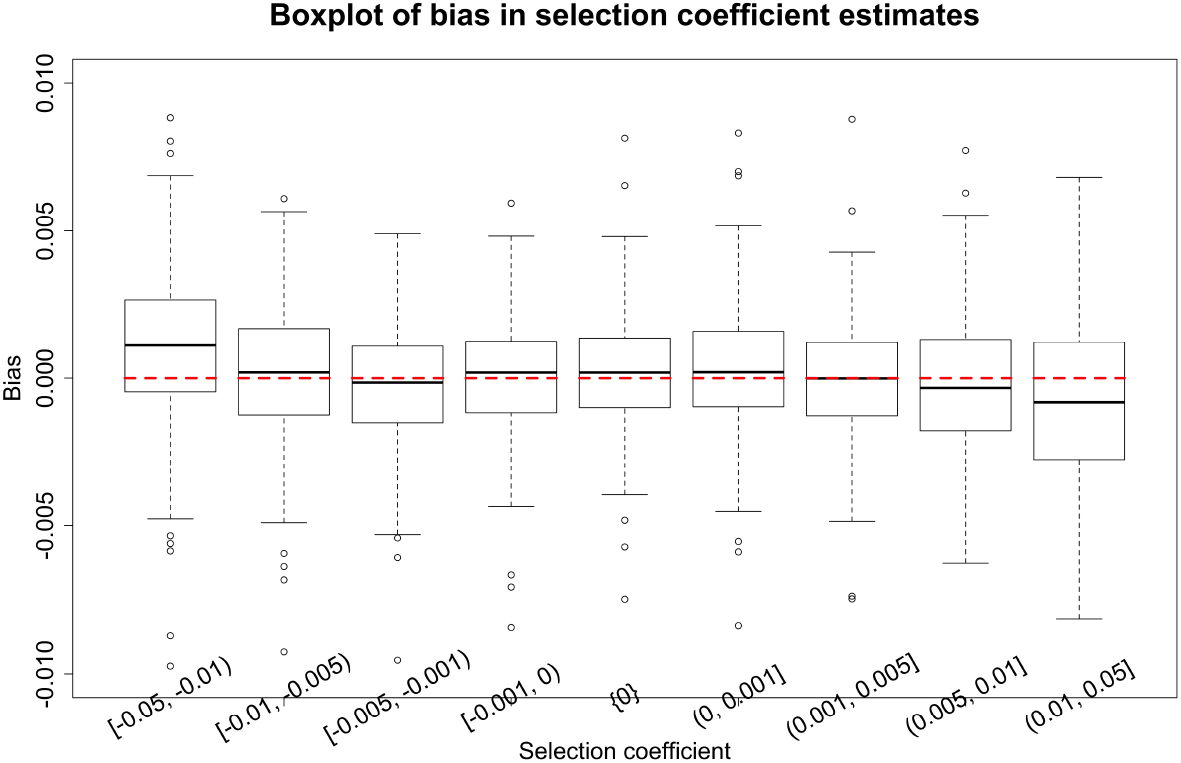
Empirical distributions for the bias in MAP estimates of the selection coefficient across different ranges of the selection coefficient *s* with the parameters *ϕ* = 0.85 and *ψ* = 1 (*i.e*., scenario D in Table 1).

We observe from Figure 4 that our estimates are nearly median-unbiased for weak selection, but the bias gets larger with an increase in the strength of selection (*i.e*., |*s*|). This bias could be caused by ascertainment arising from the procedure that we use to generate simulated datasets. In our simulation studies, only the simulated datasets in which no fixation event has occurred in the underlying population are kept. This setting means that the Wright-Fisher model in our data generation process is equivalent to that conditioned on no fixation event occurred, which does not match that in our approach for estimating the selection coefficient. Such a mismatch could bring about the underestimation of the selection coefficient (*i.e*., the simulated datasets that are retained correspond to a biased sample of the underlying population mutant allele frequency trajectories that reach loss or fixation more slowly, and therefore the estimates for the strength of selection are more likely to be smaller than their true values), especially for strong selection, since fixation events are more likely (see the histograms of the loss probability of mutations and those of the fixation probability of mutations in Supplementary Material, Figure S1 for each level of selection coefficient, where the loss probability and fixation probability are calculated with 1000 replicates for each simulated dataset). We find that the simulated datasets with large loss probabilities are all generated with the selection coefficient *s* ∈ [−0.05, −0.01) whereas those with large fixation probabilities are all generated with the selection coefficient *s* ∈ (0.01, 0.05], which correspond to the levels of selection coefficient with significant bias in Figure 4. The bias resulting from the mismatch can be fully eliminated by conditioning the Wright-Fisher diffusion to survive (He et al., 2020b).

### 3.1.2. Performance in testing selection changes

To evaluate how our approach performs for testing selection changes, we produce the receiver operating characteristic (ROC) curves across different data qualities in Figure 5, where the true-positive rate (TPR) and false-positive rate (FPR) are computed for each value of the posterior probability for the selection change that is used as a threshold to classify a locus as experiencing a shift in selection, and the ROC curve is produced by plotting the TPR against the FPR. We compute the area under the ROC curve (AUC) to summarise the performance. From Figure 5, we see that even though data qualities vary across different scenarios, all curves are very concave and close to the upper left corner of the ROC space with their AUC values varying from 0.89 to 0.94. This suggests that our procedure has superior performance in testing selection changes even for the datasets of up to around 31.0% missing rate and 13.9% error rate (see scenario A). Although these ROC curves almost overlap with each other, we can still see that improved data quality yields better performance (see, *e.g*., scenario F).

**Figure 5:**
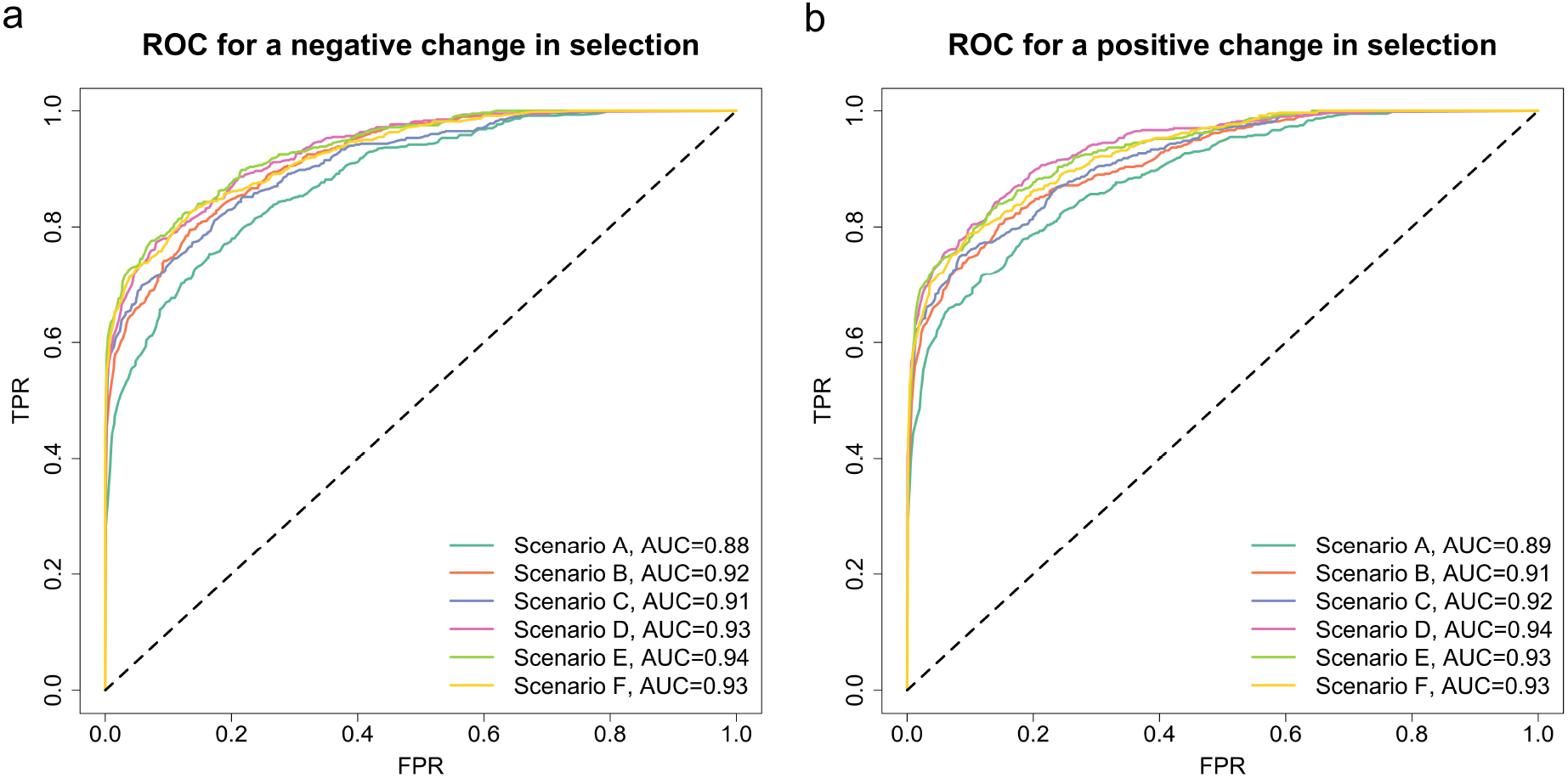
ROC curves for testing a change in selection across different data qualities and selection scenarios. The AUC value for each curve is summarised. Data qualities (scenarios A–F) are described in Table 1, and selection scenarios (scenarios 1–9) are described in Table 2. ROC curves for (a) a negative change in selection and (b) a positive change in selection.

To assess how our approach performs for different levels of selection change, in particular for small changes, we run an additional group of simulations with an example of codominance, where we adopt the parameters *ϕ* = 0.85 and *ψ* = 1. We vary the selection change Δ*s* ∈ [−0.05, 0.05], which is partitioned into nine subintervals [−0.05, −0.01), [−0.01, −0.005), [−0.005, −0.001), [−0.001, 0), {0}, (0, 0.001], (0.001, 0.005], (0.005, 0.01] and (0.01, 0.05]. For each subinterval, *e.g*., [−0.01, −0.005), we uniformly draw the selection change Δ*s* from [−0.01, −0.005), with which we uniformly pick the selection coefficient *s*^−^ from [max{−0.05, −0.05 − Δ*s*}, min{0.05, 0.05 − Δ*s*}] and thus the selection coefficient *s*^+^ = *s*^−^ + Δ*s*. We generate a dataset of genotype likelihoods with the selection coefficients *s*^−^ and *s*^+^. We repeat this procedure until we obtain 200 simulated datasets for each subinterval. Similarly, we run the ROC analysis, and the resulting ROC curves for different levels of selection change are shown in Figure 6 with their AUC values.

**Figure 6:**
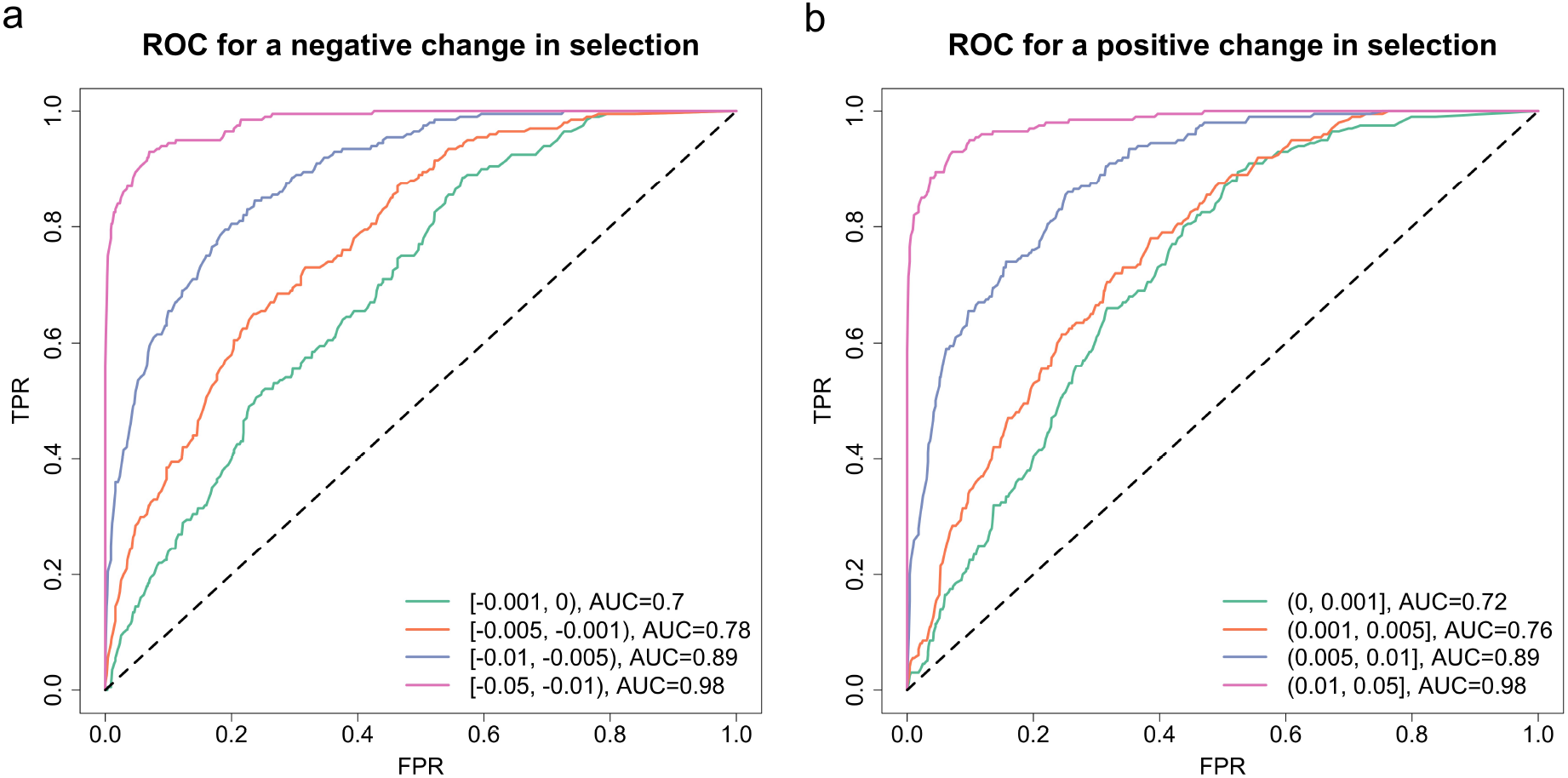
ROC curves for testing a change in selection across different ranges of the selection change Δ*s* with the parameters *ϕ* = 0.85 and *ψ* = 1 (*i.e*., scenario D in Table 1). The AUC value for each curve is summarised. ROC curves for (a) a negative change in selection and (b) a positive change in selection.

By examining the plots of the ROC curves in Figure 6, we find that the performance becomes significantly better with the increase in the degree of the change in the selection coefficient (*i.e*., |Δ*s*|) as expected. Even for small changes |Δ*s*| *<* 0.001, the AUC value is still larger than 0.70, and for large changes |Δ*s*| *>* 0.01, the AUC value is up to 0.98. Our results illustrate that our method has strong discriminating power of testing selection changes, even though such a change is small.

### 3.2. Horse coat colouration

We employ our approach to infer selection acting on the *ASIP* and *MC1R* genes associated with base coat colours (black and chestnut) and the *KIT13* and *TRPM1* genes associated with white coat patterns (tobiano and leopard complex) based on the data on aDNA sequences from previous studies of Ludwig et al. (2009), Pruvost et al. (2011) and Wutke et al. (2016), which were found to be involved in ecological, environmental and cultural shifts (Ludwig et al., 2009, 2015; Wutke et al., 2016). Since only called genotypes are available in Wutke et al. (2016), we use the following procedure to generate genotype likelihoods for each gene in the same format as those produced by GATK (McKenna et al., 2010): if the genotype is called, we set the genotype likelihood of the called genotype to 1 and those of the other two to 0, and otherwise, all possible (ordered) genotypes are assigned equal genotype likelihoods that are normalised to sum to 1. See genotype likelihoods for each gene in Supplementary Material, Table S4.

Due to the underlying assumption of our approach that mutation occurred before the initial sampling time point, in our following analysis we exclude the samples drawn before the sampling time point that the mutant allele was first found in the sample for each gene. We adopt the horse demographic history estimated by Der Sarkissian et al. (2015) (see Supplementary Material, Figure S2) with the average length of a single generation of the horse being eight years and set the reference population size *N*_0_ = 16000 (*i.e*., the most recent size of the horse population). For each gene, we take the dominance parameter *h* to be the value given in Wutke et al. (2016).

In our PMMH procedure, we run 20000 iterations with a burn-in of 10000 iterations, and the other settings are the same as we adopted in Section 3.1, including those in the Euler-Maruyama approach. The estimates of the selection coefficient and its change for each gene with their 95% HPD intervals are summarised in Supplementary Material, Table S5.

#### 3.2.1. Selection of horse base coat colours

Base coat colours in horses are primarily determined by *ASIP* and *MC1R*, which direct the type of pigment produced, black eumelanin (*ASIP*) or red pheomelanin (*MC1R*) (Corbin et al., 2020). More specifically, *ASIP* on chromosome 22 is associated with the recessive black coat, and *MC1R* on chromosome 3 is associated with the recessive chestnut coat. Ludwig et al. (2009) found that there was a rapid increase in base coat colour variation during horse domestication (starting from approximately 3500 BC) and provided strong evidence of positive selection acting on *ASIP* and *MC1R*. Fang et al. (2009) suggested that such an increase was directly caused by human preferences and demands. We apply our approach to test their hypothesis that selection acting on *ASIP* and *MC1R* was changed when horse became domesticated and estimate their selection intensities. The resulting posteriors for *ASIP* and *MC1R* are shown in Figures 7 and 8, respectively.

**Figure 7:**
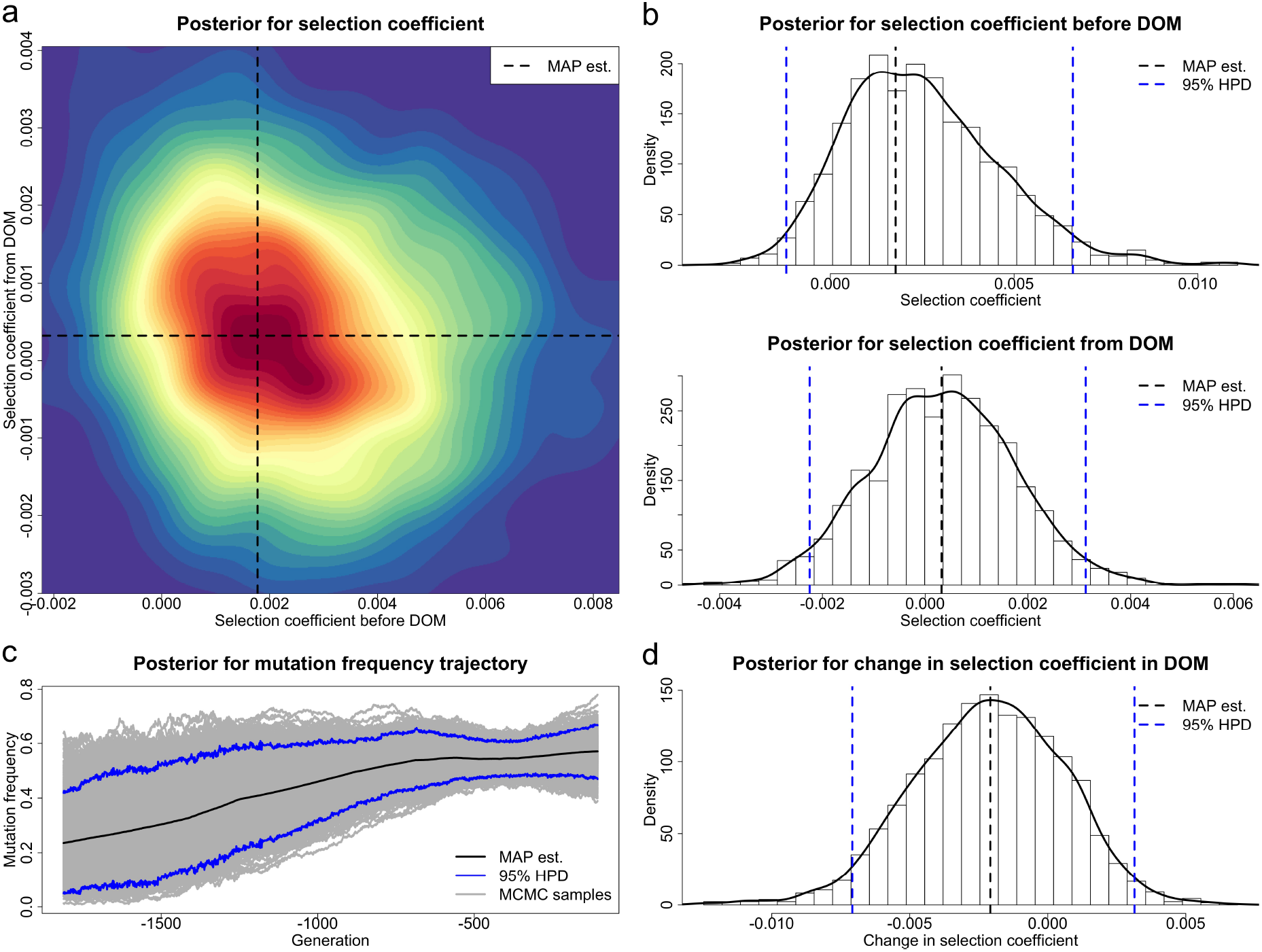
Posteriors for the selection coefficients of the *ASIP* mutation before and from horse domestication (starting from 3500 BC) and the underlying frequency trajectory of the *ASIP* mutation in the population. The samples drawn before 12500 BC are excluded. DOM stands for domestication. (a) Joint posterior for the selection coefficients *s*^−^ and *s*^+^. (b) Marginal posteriors for the selection coefficients *s*^−^ and *s*^+^. (c) Posterior for the underlying frequency trajectory of the *ASIP* mutation. (d) Posterior for the selection change Δ*s*.

**Figure 8:**
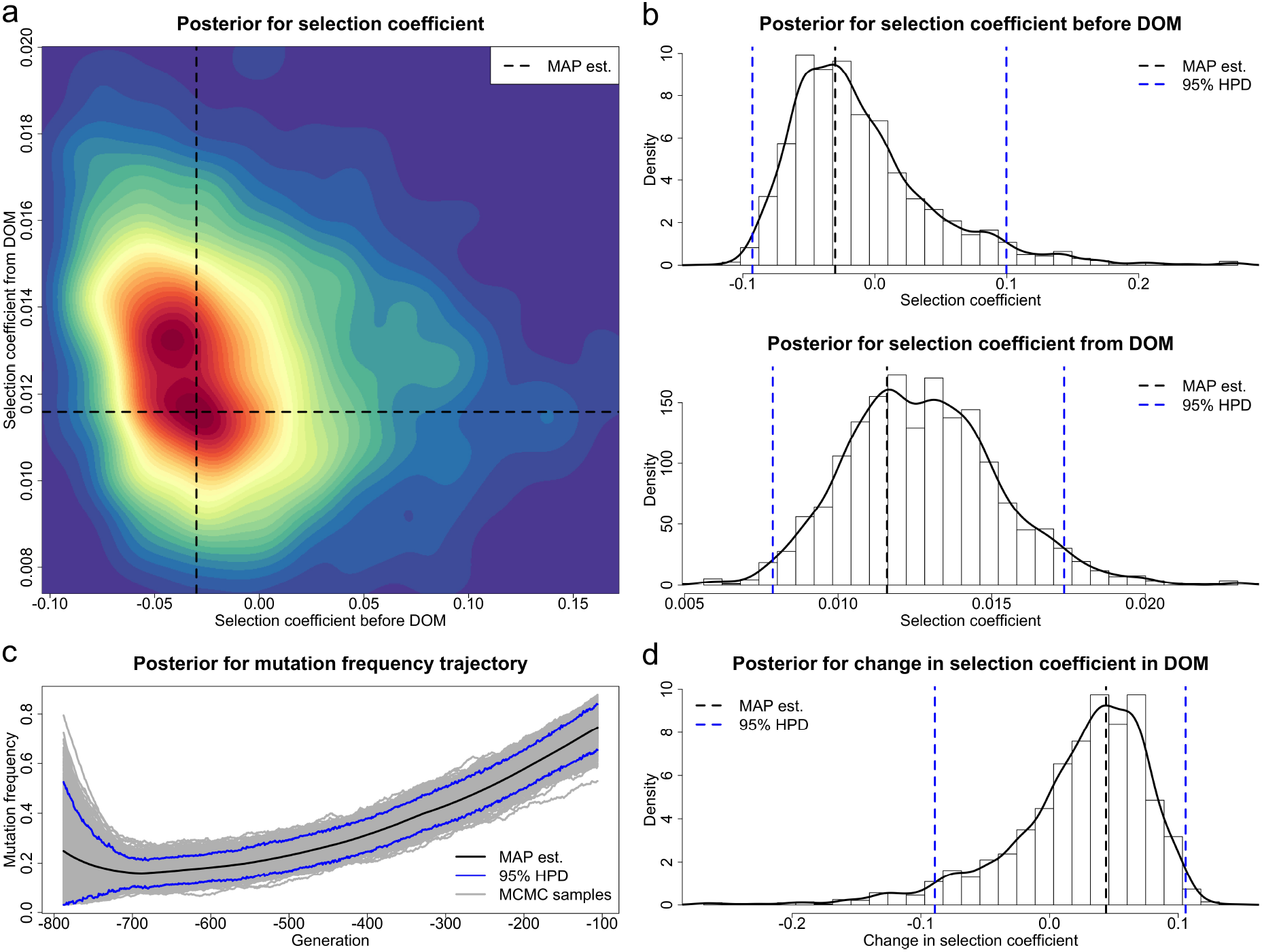
Posteriors for the selection coefficients of the *MC1R* mutation before and from horse domestication (starting from 3500 BC) and the underlying frequency trajectory of the *MC1R* mutation in the population. The samples drawn before 4300 BC are excluded. DOM stands for domestication. (a) Joint posterior for the selection coefficients *s*^−^ and *s*^+^. (b) Marginal posteriors for the selection coefficients *s*^−^ and *s*^+^. (c) Posterior for the underlying frequency trajectory of the *MC1R* mutation. (d) Posterior for the selection change Δ*s*.

Our estimate of the selection coefficient for the *ASIP* mutation is 0.0018 with 95% HPD interval [−0.0012, 0.0066] before domestication and 0.0003 with 95% HPD interval [−0.0022, 0.0031] after horses were domesticated. The 95% HPD interval contains 0 for the selection coefficient *s*^−^, but there is still some evidence showing that the *ASIP* mutation was most probably favoured by selection before domestication since the posterior probability for positive selection is 0.904. The posterior for the selection coefficient *s*^+^ is approximately symmetric about 0, which indicates that the *ASIP* mutation was effectively neutral after horse domestication started. Our estimate of the change in the selection acting on the *ASIP* mutation when horses became domesticated is −0.0021 with 95% HPD interval [−0.0071, 0.0031], and the posterior probability for such a negative change is 0.779. Our estimate for the underlying *ASIP* mutation frequency trajectory illustrates that the *ASIP* mutation frequency rises substantially in the pre-domestication period and then keeps approximately constant in the post-domestication period.

Our estimate of the selection coefficient for the *MC1R* mutation is −0.0300 with 95% HPD interval [−0.0928, 0.0997] before horses became domesticated and 0.0116 with 95% HPD interval [0.0079, 0.0174] after domestication started. Our estimates reveal that the *MC1R* mutation was effectively neutral or selectively deleterious in the pre-domestication period (with posterior probability for negative selection being 0.670) but became positively selected after horse domestication started (with posterior probability for positive selection being 1.000). Our estimate of the change in the selection acting on the *MC1R* mutation from a pre- to a post-domestication period is 0.0439 with 95% HPD interval [−0.0891, 0.1058], and the posterior probability for such a positive change is 0.748. We see a slow decline in the *MC1R* mutation frequency before domestication (even though the evidence of negative selection before domestication is weak) and then a significant increase after horse domestication started in our estimate for the underlying *MC1R* mutation frequency trajectory.

#### 3.2.2. Selection of horse white coat patterns

Tobiano is a white spotting pattern in horses characterised by patches of white that typically cross the topline somewhere between the ears and tail. It is inherited as an autosomal dominant trait that was reported in Brooks et al. (2007) to be associated with a locus in intron 13 of the *KIT* gene on chromosome 3. Wutke et al. (2016) observed that spotted coats in early domestic horses revealed a remarkable increase, but medieval horses carried significantly fewer alleles for these traits, which could result from the shift in human preferences and demands. We apply our method to test their hypothesis that selection acting on *KIT13* was changed when the medieval period began (in around AD 400) and estimate their selection intensities. We show the resulting posteriors for *KIT13* in Figure 9.

**Figure 9:**
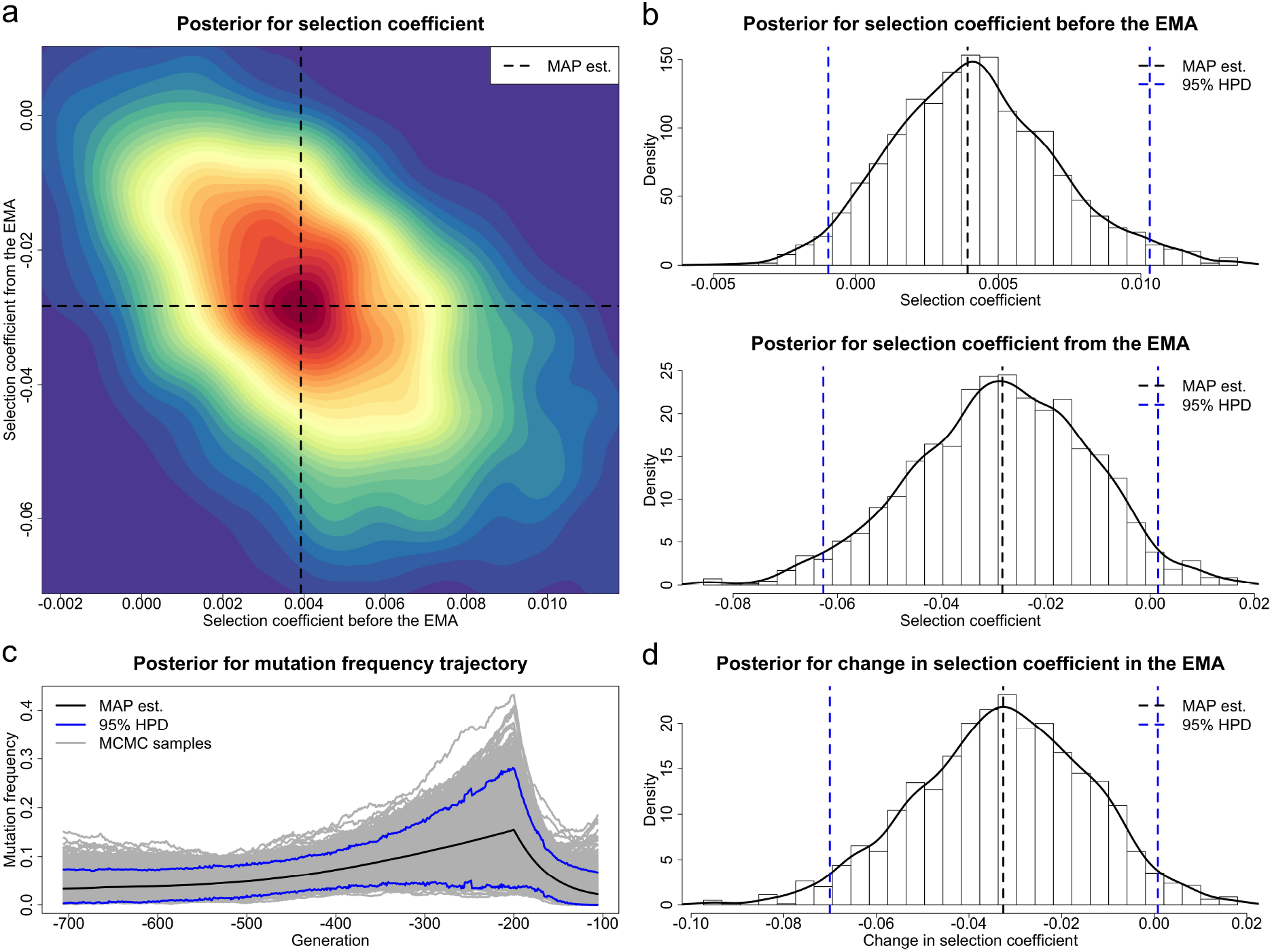
Posteriors for the selection coefficients of the *KIT13* mutation before and from the Middle Ages (starting from AD 400) and the underlying frequency trajectory of the *KIT13* mutation in the population. The samples drawn before 3645 BC are excluded. EMA stands for Early Middle Ages. (a) Joint posterior for the selection coefficients *s*^−^ and *s*^+^. (b) Marginal posteriors for the selection coefficients *s*^−^ and *s*^+^. (c) Posterior for the underlying frequency trajectory of the *KIT13* mutation. (d) Posterior for the selection change Δ*s*.

Our estimate of the selection coefficient for the *KIT13* mutation is 0.0039 with 95% HPD interval [−0.0010, 0.0103] before the medieval period, which shows that the *KIT13* mutation was positively selected before the Middle Ages (*i.e*., the posterior probability for positive selection is 0.935). Our estimate of the selection coefficient for the *KIT13* mutation is −0.0284 with 95% HPD interval [−0.0627, 0.0015] during the medieval period, which demonstrates that the *KIT13* mutation became selectively deleterious during the Middle Ages (*i.e*., the posterior probability for negative selection is 0.969). Our estimate of the change in the selection acting on the *KIT13* mutation when the Middle Ages started is −0.0326 with 95% HPD interval [−0.0700, 0.0008], and the posterior probability for such a negative change is 0.969. We observe from our estimate for the underlying *KIT13* mutation frequency trajectory that the *KIT13* mutation experienced a gradual increase after horse were domesticated and then a marked decline in the Middle Ages.

Leopard complex is a group of white spotting patterns in horses characterised by a variable amount of white in the coat with or without pigmented leopard spots, which is inherited by the incompletely dominant *TRPM1* gene residing on chromosome 1 (Terry et al., 2004). The first genetic evidence of the leopard complex coat pattern could date back to the Pleistocene (Ludwig et al., 2015). Ludwig et al. (2015) found shifts in the selection pressure for the leopard complex coat pattern in domestic horses but did not investigate whether *TRPM1* undergone a change in selection from a pre- to a post-domestication period. We apply our method to test the hypothesis that selection acting on *TRPM1* was changed when horses became domesticated and estimate their selection intensities. The resulting posteriors for *TRPM1* are shown in Figure 10. Our estimate of the selection coefficient for the *TRPM1* mutation is −0.0005 with 95% HPD interval [−0.0042, 0.0029] before horses were domesticated and −0.0078 with 95% HPD interval [−0.0139, −0.0005] after domestication started. Our estimates provide little evidence of negative selection in the pre-domestication period (with posterior probability for negative selection being 0.617) but strong evidence of negative selection in the post-domestication period (with posterior probability for negative selection being 0.980). Our estimate of the change in the selection acting on the *TRPM1* mutation when horses became domesticated is −0.0066 with 95% HPD interval [−0.0142, 0.0025]. The 95% HPD interval for the change in the selection coefficient contains 0, which however still provides sufficient evidence to support that a negative change took place in selection when horses became domesticated (*i.e*., the posterior probability for such a negative change is 0.942). Our estimate for the underlying trajectory of the *TRPM1* mutation frequency displays a slow decrease in the *TRPM1* mutation frequency during the pre-domestication period with a significant drop after horse domestication started.

**Figure 10:**
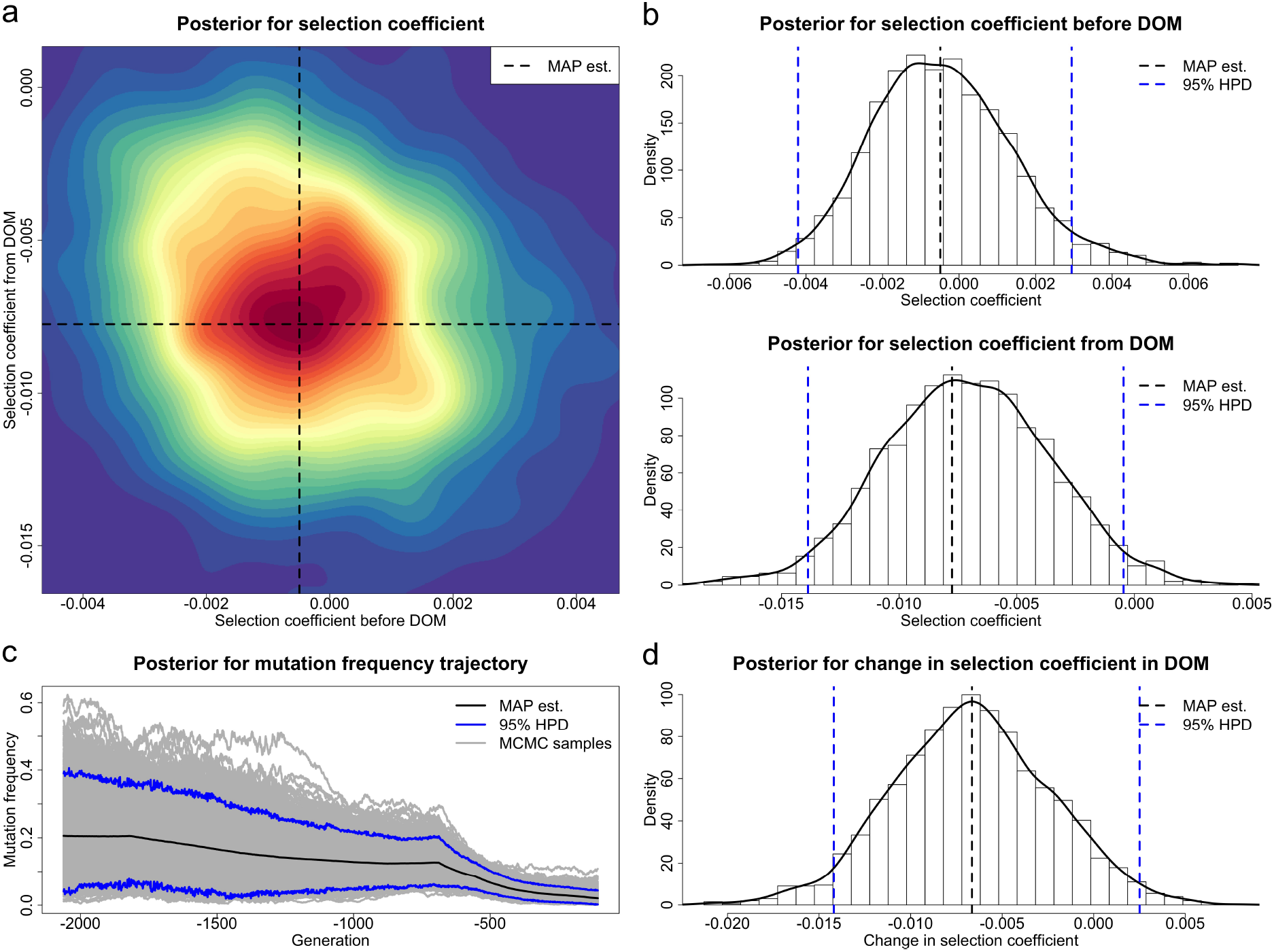
Posteriors for the selection coefficients of the *TRPM1* mutation before and from horse domestication (starting from 3500 BC) and the underlying frequency trajectory of the *TRPM1* mutation in the population. The samples drawn before 14500 BC are excluded. DOM stands for domestication. (a) Joint posterior for the selection coefficients *s*^−^ and *s*^+^. (b) Marginal posteriors for the selection coefficients *s*^−^ and *s*^+^. (c) Posterior for the underlying frequency trajectory of the *TRPM1* mutation. (d) Posterior for the selection change Δ*s*.

## 4. Discussion

In this work, we have introduced a novel Bayesian approach for estimating temporally changing selection intensity from aDNA data. To our knowledge, most earlier methods ignore sample uncertainties resulting from the damage and fragmentation of aDNA molecules, which however are the main characteristics of aDNA. Our method to circumvent this issue is that we base our selection inference on genotype likelihoods rather than called genotypes, which facilitates the incorporation of genotype uncertainties. We have developed a novel two-layer HMM framework, where the top hidden layer models the underlying frequency trajectory of the mutant allele in the population, the intermediate hidden layer models the unobserved genotype of the individual in the sample, and the bottom observed layer denotes the data on aDNA sequences. By working with the PMMH algorithm, where the marginal likelihood is approximated through particle filtering, our approach enables us to reconstruct the underlying population allele frequency trajectories. Moreover, our method provides the flexibility of modelling time-varying demographic histories.

The performance of our approach has been evaluated through extensive simulations, which show that our procedure can produce accurate selection inference from aDNA data across different evolutionary scenarios even though samples are sparsely distributed in time with small sizes and in poor quality. Note that in our simulations studies, our procedure has only been tested in the case of a bottleneck demographic history and codominance, but in principle, the conclusions we have drawn here hold for other demographic histories and levels of dominance. Demographic histories have been demonstrated to have little influence on the inference of selection from time series genetic data in Jewett et al. (2016), but in practice, demographic history misspecification can largely bias the inference of selection (see, *e.g*., Schraiber et al., 2016). We leave additional simulation studies for the scenarios of the mutant allele being recessive (*h* = 0) and dominant (*h* = 1) in Supplementary Material, Figures S3 and S4, respectively, as well as in Table S6.

We have illustrated the utility of our approach with an application to ancient horse samples from earlier studies of Ludwig et al. (2009), Pruvost et al. (2011) and Wutke et al. (2016), which were genotyped at the loci (*e.g*., *ASIP, MC1R, KIT13* and *TRPM1*) for horse coat colouration. The published demographic estimate for the horse population from Der Sarkissian et al. (2015) is utilised in our analysis. Our findings are consistent with previous studies that the coat colour variation in the horse is a domestic trait that was subject to early selection by humans (Hunter, 2018), *e.g*., *ASIP, MC1R* and *TRPM1*, and human preferences have changed greatly over time and across cultures (Wutke et al., 2016), *e.g*., *KIT13*. We have also run the analysis with a fixed demographic history of *N* = 16000 (*i.e*., the most recent size of the horse population estimated by Der Sarkissian et al. (2015)), which produces similar results (see Supplementary Material, Figures S5–S8 and Table S7).

Our results for the base coat colour are consistent with previous studies that the shift in horse coat colour variation in the early stage of domestication could be caused by relaxed selection for camouflage alleles (Hunter, 2018). More specifically, the *ASIP* mutation was positively selected in the pre-domestication period, but the *MC1R* mutation was not. From Sandoval-Castellanos et al. (2017), forest cover was growing as a result of global warming during the Late Pleistocene, which pushed horses into the forest full of predators. Dark-coloured coats could help horses avoid predators through better camouflage, therefore improving their chances of survival. After horses were domesticated, the *ASIP* mutation was no longer selectively advantageous, but the *MC1R* mutation became favoured by selection. The shift in the horse coat colour preference from dark to light could be explained by that light-coloured horses were no longer required to be protected against predation due to domestication. Furthermore, light-coloured coats could facilitate horse husbandry since it was easier to keep track of the horses that were not camouflaged (Fang et al., 2009).

Our results for the tobiano coat pattern illustrate that the *KIT13* mutation was favoured by selection from domestication till the Middle Ages and then became negatively selected, which confirm the findings of Wutke et al. (2016). Such a negative change in selection of the tobiano coat could result from pleiotropic disadvantages, a lower religious prestige, a reduced need to separate domestic horses from their wild counterparts or novel developments in weaponry during the medieval period (see Wutke et al., 2016, and references therein).

Our results for the leopard complex coat pattern demonstrate that the *TRPM1* mutation was negatively selected from the Late Pleistocene onwards. Our evidence of negative selection acting on the *TRPM1* mutation in the pre-domestication period is not strong enough, but we can still see a slow drop in the *TRPM1* mutation frequency over time. The *TRPM1* mutation is the most common cause of congenital stationary night blindness (CSNB) (Bellone et al., 2013), which could reduce the chance to survive in the wild since vision is key for communication, localisation, orientation, avoiding predators and looking for food (Murphy et al., 2009). The weak intensity of negative selection could be explained as resulting in part from that only horses homozygous for the leopard complex coat pattern are influenced by CSNB, which however remarkably increased when horses were domesticated. In the post-domestication period, horses were harnessed mainly for power and transportation, *e.g*., they were used to pull wheeled vehicles, chariots, carts and wagons in the early stage of domestication and later used in war, in hunting and as a means of transport, which all strongly rely on the ability to see. Moreover, night-blind horses are nervous and timid in human care, and difficult to handle at dusk and darkness (Rebhun et al., 1984).

These genes have been well studied by other methods (*e.g*., Bollback et al., 2008; Malaspinas et al., 2012; Steinrücken et al., 2014; Schraiber et al., 2016; He et al., 2020a,b), but our results are not completely consistent with those presented in previous studies (see Supplementary Material, Table S8 for a summary of the results produced by existing approaches). The discrepancy can be mainly explained by that the strength of selection is assumed to be fixed over time in all existing methods. Other potential causes of the discrepancy are *e.g*., only 89 ancient horse samples from Ludwig et al. (2009) being used to infer selection acting on the *ASIP* and *MC1R* mutations (*e.g*., Malaspinas et al., 2012; Steinrücken et al., 2014; Schraiber et al., 2016) and a more adequate modelling of relevant scenarios (*e.g*., He et al. (2020a) modelled genetic recombination and local linkage in the inference of selection acting on the *KIT13* and *KIT16* mutations).

In aDNA studies, the samples with missing genotypes are usually filtered, and the remaining samples are manually grouped into a small number of sampling time points, *e.g*., ancient horse samples were grouped into six sampling time points in Ludwig et al. (2009) or nine sampling time points in Wutke et al. (2016). Grouping has been shown to significantly alter the results of the inference of selection from time series genetic data in He et al. (2020b). Our approach provides an alternative that can address the issue caused by the procedure of sample filtering and grouping (see Supplementary Material, Figures S9 and S10, as well as Table S9, for additional simulation studies, where we compare our results based on genotype likelihoods with those produced with called genotypes). Our simulation studies show that the commonly used procedure of processing aDNA data for the inference of selection can significantly bias the result, in particular for poor data quality, and suggest that the method based on genotype likelihoods can be a more promising alternative for future aDNA studies.

In our work, the demographic history is required to be prespecified but allowed to be changed over time, as in Schraiber et al. (2016) and He et al. (2020b). Our method is ready to be extended to co-estimate the population size like Malaspinas et al. (2012), but genetic variation at a single locus is not sufficient to produce a reliable estimate of the population size, *e.g*., the population size estimated by Malaspinas et al. (2012) from allele frequency time series data at *ASIP* is far smaller than the genome-wide estimate produced by Der Sarkissian et al. (2015), and therefore Malaspinas et al. (2012) could not distinguish positive from negative selection for *ASIP*, which has been shown to be positively selected in other studies (*e.g*., Ludwig et al., 2009; Steinrücken et al., 2014; Schraiber et al., 2016; He et al., 2020b). Several methods like Foll et al. (2015) and Ferrer-Admetlla et al. (2016) can jointly estimate the population size and selection coefficients from time series genomic data but are usually used in experimental evolution rather than aDNA studies due to computational efficiency. Moreover, most of these methods are commonly divided into two steps (but see Ferrer-Admetlla et al., 2016). The first step is to estimate the population size from all loci across the genome assuming selective neutrality, and in the second step, given the estimate of the population size, the selection coefficient for each single locus is independently inferred. These two-step procedures suffer from the difficulty of rejecting the null hypothesis of selective neutrality and the issue of underestimating the population size and selection coefficients (Paris et al., 2019), in particular with an increase in the proportion of selected loci across the genome. A method to address this issue can be to co-estimate the population size with purifying and background selection in the first step as a null model and then fix that model in the second step (Johri et al., 2020, 2021, 2022a,b). Joint inference of demographic and selective parameters from ancient genomes, an important topic of future investigation, is anticipated to significantly improve the estimation of selection coefficients and even permit the inference of demographic changes.

Although the level of dominance is prespecified in our analysis of simulated data and aDNA data such as Malaspinas et al. (2012) and He et al. (2020a,b), our method allows to co-estimate the dominance parameter (see Supplementary Material, Figures S11 and S12, as well as Table S10, for additional simulation studies evaluating its performance in co-estimating the dominance parameter). In practice, a reliable estimate of the dominance parameter depends heavily on the quality and quantity of data, *e.g*., Foll et al. (2015) illustrated that WFABC performed well in the joint estimation of the dominance parameter from the *Panaxia dominula* data (*i.e*., a total of 58592 samples distributed over 60 generations, 51 sampling time points) but exhibited poor performance in the simulation studies of Kojima et al. (2020), where three replicated populations were simulated, each with 250 samples distributed over 60 generations, five sampling time points. Considering that the quality and quantity of aDNA data are poor, prespecifying the dominance parameter in aDNA studies with sufficient prior knowledge can be a feasible and reasonable alternative.

Compared to existing methods (*e.g*., Bollback et al., 2008; Malaspinas et al., 2012; Steinrücken et al., 2014; Schraiber et al., 2016; Ferrer-Admetlla et al., 2016; He et al., 2020a,b), our Bayesian procedure enables the selection coefficient to vary in time (*i.e*., piecewise constant) although the event that might change selection is required to be prespecified. However, this is still important in aDNA studies as adaptation in natural populations often involves adaptation to ecological, environmental and cultural shifts. We run our procedure on the ancient horse samples presented in Supplementary Material, Table S4 with the same settings as we adopted in Section 3.2, except that the selection coefficient is fixed over time (see Supplementary Material, Figure S13 for the resulting posteriors and Table S11 for the estimates of the selection coefficients with their 95% HPD intervals). We find for example that the *KIT13* mutation was effectively neutral during the post-domestication period, which contradicts the archaeological evidence and historical records that spotted horses were subject to early selection by humans, but the preference shifted in the medieval period (see Wutke et al., 2016, and references therein). Compared to the results shown in Figures 7–10, we see the necessity of modelling temporally variable selection in aDNA studies. Our procedure lends itself to being extended to allow multiple events that might change selection (see Section 5.2). To guarantee computational efficiency, a feasible solution is to adopt an adaptive strategy that allows for automatically tuning the selection coefficients during a run (see Luengo et al., 2020, for a review). A potential direction for future research is the inference of selection and its strength and timing of changes from time serial genetic samples (Shim et al., 2016; Mathieson, 2020).

One fundamental limitation of our method applied for the inference of selection from aDNA data is that it assumes that all samples have been collected after the mutant allele was created. However, allele age is not always available, and as a result, in our analysis we have to exclude the samples drawn before the time that the mutant allele was first observed in the sample, which might alter the result of the inference of selection. To address this problem, we can extend our approach to jointly estimate the allele age as in Malaspinas et al. (2012), Schraiber et al. (2016) and He et al. (2020b), and the foreseeable challenge is how to resolve particle degeneracy and impoverishment issues in our PMMH-based procedure that result from low-frequency mutant alleles at the early stage facing a higher probability of being lost. Also, similar to most existing approaches (except for Mathieson & McVean (2013) and Lyu et al. (2022)), our method lacks the ability to take gene migration into account, which is a common source of confounding in aDNA studies of adaptive processes since natural populations are almost always structured (Mathieson et al., 2015). To resolve this issue, we can incorporate migration into the Wright-Fisher model with selection and co-estimate the migration rate through our PMMH-based procedure, where following Lyu et al. (2022), we alternatively update the selection-related and migration-related parameters to improve the mixing of the chain. In addition, our method assumes that the gene of our interest is independent of others, which however can be easily violated once there exist interactions between genes like epistasis and linkage, *e.g*., *MC1R* is epistatic to *MC1R* (Rieder et al., 2001), and *KIT13* is tightly linked to *KIT16* (Dumont & Payseur, 2008). Ignoring gene interactions might bias the inference of selection (He et al., 2020a). To extend our method to the scenario of multiple genes with epistasis and/or linkage, we need to find a good approximation for the Wright-Fisher model of multiple genes evolving under selection with epistasis and linkage over time, which becomes challenging with an increase in the number of genes. We will discuss how to extend our method to the case of two genes with epistasis and/or linkage in our upcoming work, and the extension for the scenario of multiple interacting genes (≥ 3) will be the topic of future investigation.

## 5. Materials and Methods

In this section, we describe in detail our approach to infer temporally variable selection from aDNA data while modelling sample uncertainties resulting from the damage and fragmentation of aDNA molecules.

### 5.1. Wright-Fisher diffusion

Let us consider a diploid population of *N* randomly mating individuals at a single locus 𝒜, which evolves subject to selection under the Wright-Fisher model (see, *e.g*., Durrett, 2008). We assume discrete time, non-overlapping generations and non-constant population size. Suppose that at locus 𝒜 there are two possible allele types, labelled 𝒜_0_ and 𝒜_1_, respectively. We attach the symbol 𝒜 _0_ to the ancestral allele, which originally exists in the population, and the symbol 𝒜 _1_ to the mutant allele, which arises in the population only once. We let selection take the form of viability selection and take per-generation relative viabilities of the three possible genotypes 𝒜_0_𝒜_0_, 𝒜_0_𝒜_1_ and 𝒜_1_𝒜 _1_ to be 1, 1 + *hs* and 1 + *s*, respectively, where *s* ∈ [−1, +∞) is the selection coefficient and *h* ∈ [0, 1] is the dominance parameter.

We now consider the standard diffusion limit of the Wright-Fisher model with selection. We measure time in units of 2*N*_0_ generations, denoted by *t*, where *N*_0_ is an arbitrary reference population size fixed through time, and assume that the population size changes deterministically, with *N* (*t*) being the number of diploid individuals in the population at time *t*. In the diffusion limit of the Wright-Fisher model with selection, as the reference population size *N*_0_ goes to infinity, the scaled selection coefficient *α* = 2*N*_0_*s* is kept constant and the ratio of the population size to the reference population size *N* (*t*)*/N*_0_ converges to a function *β*(*t*). As demonstrated in Durrett (2008), the mutant allele frequency trajectory through time converges to the diffusion limit of the Wright-Fisher model with the reference population size *N*_0_ approaching infinity. We refer to this diffusion limit as the Wright-Fisher diffusion with selection.

We let *X* denote the Wright-Fisher diffusion with selection, which models the mutant allele frequency evolving in the state space [0, 1] under selection. Many existing approaches define the Wright-Fisher diffusion *X* in terms of the partial differential equation (PDE) that characterises its transition probability density function (*e.g*., Bollback et al., 2008; Steinrücken et al., 2014; He et al., 2020b). Instead, like Schraiber et al. (2016), He et al. (2020a) and Lyu et al. (2022), we define the Wright-Fisher diffusion *X* as the solution to the stochastic differential equation

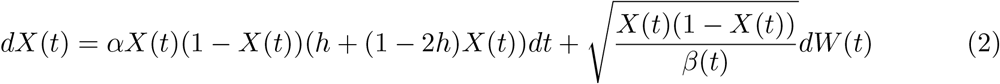

for *t* ≥ *t*_0_ with initial condition *X*(*t*_0_) = *x*_0_, where *W* represents the standard Wiener process.

### 5.2. Bayesian inference of selection

Suppose that the available data are always drawn from the underlying population at a finite number of distinct time points, say *t*_1_ *< t*_2_ *<* … *< t*_*K*_, measured in units of 2*N*_0_ generations. At the sampling time point *t*_*k*_, there are *N*_*k*_ individuals sampled from the underlying population, and for individual *n*, let ***r***_*n,k*_ represent, in this generic notation, all of the reads at the locus of interest. The population genetic parameters of interest in this work are the selection coefficient *s* and dominance parameter *h*, and for ease of notation, we set ***ϑ*** = (*s, h*) in what follows.

#### 5.2.1. Hidden Markov model

Our method is built upon the HMM framework introduced by Bollback et al. (2008), where the underlying population evolves under the Wright-Fisher diffusion with selection in Eq. (2), and the observed sample is made up of the individuals independently drawn from the underlying population at each sampling time point. To model sample uncertainties resulting from the damage and fragmentation of aDNA molecules, our approach infers selection from raw reads rather than called genotypes. We let ***x***_1:*K*_ = {*x*_1_, *x*_2_, …, *x*_*K*_} represent the frequency trajectory of the mutant allele in the underlying population at the sampling time points ***t***_1:*K*_, and the posterior probability distribution for the population genetic parameters ***ϑ*** and population mutant allele frequency trajectory ***x***_1:*K*_ (up to proportionality) can be formulated as

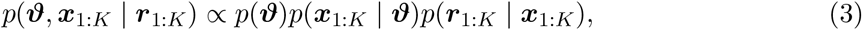

where ***r***_1:*K*_ = {***r***_1_, ***r***_2_, …, ***r***_*K*_} with 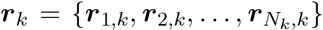. In Eq. (3), *p*(***ϑ***) is the prior probability distribution for the population genetic parameters and can be taken to be a uniform prior over the parameter space if prior knowledge is poor, *p*(***x***_1:*K*_ | ***ϑ***) is the probability distribution for the mutant allele frequency trajectory of the underlying population at all sampling time points, and *p*(***r***_1:*K*_ | ***x***_1:*K*_) is the probability of observing the reads of all sampled individuals given the mutant allele frequency trajectory of the population.

With the Markov property of the Wright-Fisher diffusion, we can decompose the probability distribution *p*(***x***_1:*K*_ | ***ϑ***) as

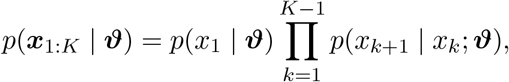

where *p*(*x*_1_ | ***ϑ***) is the prior probability distribution for the starting population mutant allele frequency, taken to be a uniform distribution over the state space [0, 1] if the prior knowledge is poor, and *p*(*x*_*k*+1_ | *x*_*k*_; ***ϑ***) is the transition probability density function of the Wright-Fisher diffusion *X* between two consecutive sampling time points for *k* = 1, 2, …, *K* − 1, satisfying the Kolmogorov backward equation (or its adjoint) resulting from the Wright-Fisher diffusion.

To calculate the probability *p*(***r***_1:*K*_ | ***x***_1:*K*_), we introduce an additional hidden layer into our HMM framework to denote the latent genotypes of all sampled individuals (see Figure 1 for the graphical representation of our two-layer HMM framework). We let ***g***_1:*K*_ = {***g***_1_, ***g***_2_, …, ***g***_*K*_} be the genotypes of the individuals drawn from the underlying population at the sampling time points ***t***_1:*K*_ with 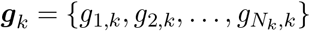, where *g*_*n,k*_ ∈ {0, 1, 2} denotes the number of mutant alleles in individual *n* at sampling time point *t*_*k*_. We then have

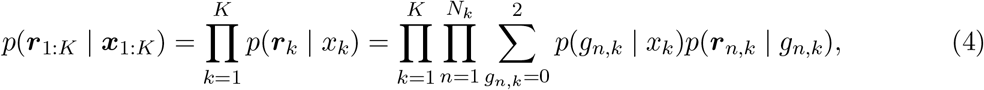

where *p*(*g*_*n,k*_ | *x*_*k*_) represents the probability distribution for the genotype *g*_*n,k*_ of individual *n* in the sample given the population mutant allele frequency *x*_*k*_, and *p*(***r***_*n,k*_ | *g*_*n,k*_) represents the probability of observing reads ***r***_*n,k*_ of individual *n* in the sample given the genotype *g*_*n,k*_, known as the genotype likelihood. Under the assumption that all individuals in the sample are drawn from the underlying population in their adulthood (*i.e*., the stage after selection but before reproduction in the life cycle, see He et al. (2017)), the first term in Eq. (4), *p*(*g*_*n,k*_ | *x*_*k*_), can be calculated through Eq. (1). Genotype likelihoods are typically calculated from aligned reads and quality scores in the process of determining the genotype for each individual through genotype calling software (*e.g*., Li et al., 2009a,b; DePristo et al., 2011), which is an essential prerequisite for most aDNA studies, therefore assuming that the second term in Eq. (4), *p*(***r***_*n,k*_ | *g*_*n,k*_), is available.

#### 5.2.2. Particle marginal Metropolis-Hastings

Since the posterior *p*(***ϑ, x***_1:*K*_ | ***r***_1:*K*_) is not available in a closed form, we resort to the PMMH algorithm introduced by Andrieu et al. (2010) in this work, which has already been successfully applied in population genetic studies (see, *e.g*., He et al., 2020b; Lyu et al., 2022). The PMMH algorithm calculates the acceptance ratio with the estimate of the marginal likelihood *p*(***r***_1:*K*_ | ***ϑ***) in the Metropolis-Hastings procedure and generates a new candidate of the population mutant allele frequency trajectory ***x***_1:*K*_ from the approximation of the smoothing distribution *p*(***x***_1:*K*_ | ***r***_1:*K*_, ***ϑ***), which can both be achieved through the bootstrap particle filter developed by Gordon et al. (1993). Such a setup permits a joint update of the population genetic parameters ***ϑ*** and population mutant allele frequency trajectory ***x***_1:*K*_.

A key ingredient of the PMMH algorithm is to construct a bootstrap particle filter that can target the smoothing distribution *p*(***x***_1:*K*_ | ***r***_1:*K*_, ***ϑ***). More specifically, at the sampling time point *t*_*k*+1_, our objective is to generate a sample from the filtering distribution *p*(*x*_*k*+1_ | ***r***_1:*k*+1_, ***ϑ***). Up to proportionality, the filtering distribution *p*(*x*_*k*+1_ | ***r***_1:*k*+1_, ***ϑ***) can be formulated as

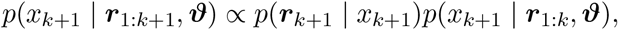

where

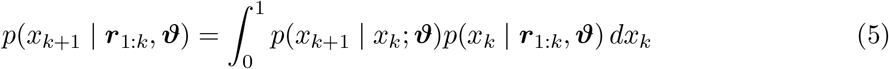

is the predictive distribution, but not available in a closed form. From Eq. (5), the predictive distribution *p*(*x*_*k*+1_ | ***r***_1:*k*_, ***ϑ***) can be approximated with a set of particles 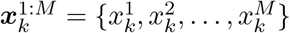 generated from the filtering distribution *p*(*x*_*k*_ | ***r***_1:*k*_, ***ϑ***) with each particle 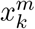 being assigned a weight 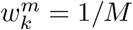, thereby

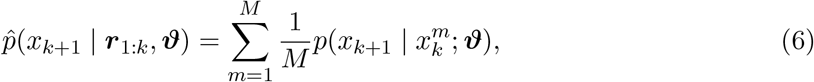

where superscript represents the particle label. By substituting Eq. (6) into Eq. (5), the filtering distribution *p*(*x*_*k*+1_ | ***r***_1:*k*+1_, ***ϑ***) (up to proportionality) can be approximated by

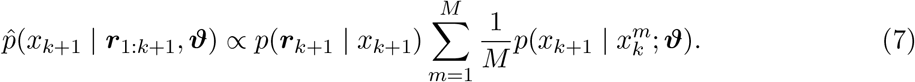

From Eq. (7), the approximation of the filtering distribution *p*(*x*_*k*+1_ | ***r***_1:*k*+1_, ***ϑ***) can be sampled with importance sampling, where we generate a set of particles 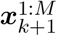 from the predictive distribution 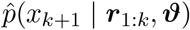 with each particle 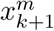 being assigned a weight 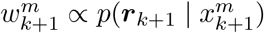 (Gordon et al., 1993). We resample *M* particles with replacement amongst the set of particles 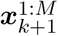 with probabilities proportional to weights 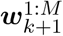. For clarity, we write down the bootstrap particle filter algorithm:

Step 1: Initialise the particles at the sampling time point *t*_1_:

Step 1a: Draw 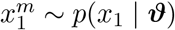 for *m* = 1, 2, …, *M*.

Step 1b: 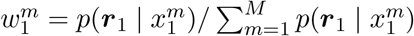 for *m* = 1, 2, …, *M*.

Step 1c: Resample *M* particles with replacement amongst 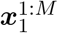 with 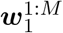.

Repeat Step 2 for *k* = 2, 3, …, *K*:

Step 2: Update the particles at the sampling time point *t*_*k*_:

Step 2a: Draw 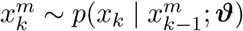 for *m* = 1, 2, …, *M*.

Step 2b: Set 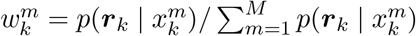 for *m* = 1, 2, …, *M*.

Step 2c: Resample *M* particles with replacement amongst 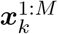 with 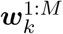.

With the procedure for the bootstrap particle filter described above, the smoothing distribution *p*(***x***_1:*K*_ | ***r***_1:*K*_, ***ϑ***) can be sampled by uniformly drawing amongst the set of particles 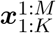, and the marginal likelihood *p*(***r***_1:*K*_ | ***ϑ***) can be estimated by

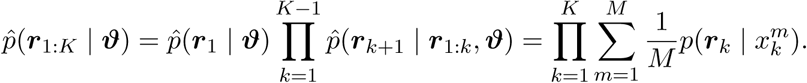

We can therefore write down the PMMH algorithm:

Step 1: Initialise the population genetic parameters ***ϑ*** and population mutant allele frequency trajectory ***x***_1:*K*_:

Step 1a: Draw ***ϑ***^1^ ∼ *p*(***ϑ***).

Step 1b: Run a bootstrap particle filter with ***ϑ***^1^ to yield 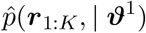 and 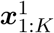.

Repeat Step 2 until a sufficient number of samples of the population genetic parameters ***ϑ*** and population mutant allele frequency trajectory ***x***_1:*K*_ have been obtained:

Step 2: Update the population genetic parameters ***ϑ*** and population mutant allele frequency trajectory ***x***_1:*K*_:

Step 2a: Draw ***ϑ***^*i*^ ∼ *q*(***ϑ*** | ***ϑ***^*i*−1^).

Step 2b: Run a bootstrap particle filter with ***ϑ***^*i*^ to yield 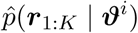 and 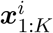.

Step 2c: Accept ***ϑ***^*i*^ and 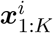 with

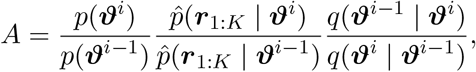

otherwise set ***ϑ***^*i*^ = ***ϑ***^*i*−1^ and 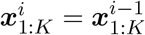.

Note that superscript represents the iteration in our procedure described above. In the sequel, we adopt random walk proposals for each component of the population genetic parameters ***ϑ*** unless otherwise specified.

Once enough samples of the population genetic parameters ***ϑ*** and population mutant allele frequency trajectory ***x***_1:*K*_ have been obtained, we can get the MAP estimates for the population genetic parameters ***ϑ*** by computing the posterior *p*(***ϑ*** | ***r***_1:*K*_) with nonparametric density estimation techniques (see Izenman, 1991, for a review). Alternatively, we can yield the minimum mean square error estimates for the population genetic parameters ***ϑ*** through their posterior means. Similarly, we can take the posterior mean of the samples of the population mutant allele frequency trajectory to be our estimate for the population mutant allele frequency trajectory ***x***_1:*K*_.

Our method allows the selection coefficient *s* to be piecewise constant over time. For example, we let the selection coefficient *s*(*t*) = *s*^−^ if *t < τ*, otherwise *s*(*t*) = *s*^+^, where *τ* is the time of an event that might change selection, *e.g*., the times of plant and animal domestication. Our procedure can then be directly applied to estimate the selection coefficients *s*^−^ and *s*^+^ for any prespecified time *τ*. The only modification required is to simulate the underlying mutant allele frequency trajectory of the population through the Wright-Fisher diffusion with the selection coefficient *s*^−^ for *t < τ* and *s*^+^ for *t* ≥ *τ*, respectively. In this case, our method also provides a procedure to estimate and test the change in the selection coefficient, denoted by Δ*s* = *s*^+^ − *s*^−^, at time *τ* by computing the posterior *p*(Δ*s* | ***r***_1:*K*_) from the samples of the selection coefficients *s*^−^ and *s*^+^. This is a highly desirable feature in aDNA studies as it enables us to test hypotheses about whether the shift in selection is linked to specific ecological, environmental and cultural drivers.

## Supplementary Material

Supplementary data are available at Molecular Biology and Evolution online.

## Acknowledgements

We thank the anonymous reviewers and the editor for their helpful comments on the earlier version of this work. This work was carried out using the computational facilities of the Advanced Computing Research Centre, University of Bristol - http://www.bristol.ac.uk/acrc/.

## Author Contributions

Z.H. designed the project and developed the method; Z.H., X.D. and W.L. implemented the method; X.D. and W.L. analysed the data under the supervision of Z.H., M.B. and F.Y.; Z.H. and X.D. wrote the manuscript; W.L., M.B. and F.Y. reviewed the manuscript.

## Data Availability

The authors state that all data necessary for confirming the conclusions of the present work are represented fully within the article. Source code implementing the approach described in this work is available at https://github.com/zhangyi-he/WFM-1L-DiffusApprox-PMMH/.

## Notes

### Competing Interest Statement

The authors have declared no competing interest.

### Summary of Updates

The Discussion section has been updated to further discuss the joint estimation of the population size and dominance parameter.

