## Supplementary Material for "Estimating temporally variable selection intensity from ancient DNA data"

| | Sel. coeff. $s^-$ | | Sel. coeff. $s^+$ | |
|---|---|---|---|---|
| Scenario | Bias | RMSE | Bias | RMSE |
| A | 0.00126 | 0.00391 | 0.00684 | 0.00952 |
| B | 0.00124 | 0.00424 | 0.00229 | 0.00708 |
| C | 0.00054 | 0.00399 | 0.00378 | 0.00806 |
| D | 0.00022 | 0.00348 | 0.00073 | 0.00595 |
| E | 0.00081 | 0.00354 | 0.00104 | 0.00603 |
| F | 0.00043 | 0.00402 | 0.00053 | 0.00618 |

(a) Scenario 1: $s^- < 0$ and $s^+ < s^-$.

| | Sel. coeff. $s^-$ | | Sel. coeff. $s^+$ | |
|---|---|---|---|---|
| Scenario | Bias | RMSE | Bias | RMSE |
| A | 0.00258 | 0.00438 | 0.00358 | 0.00646 |
| B | 0.00111 | 0.00411 | 0.00100 | 0.00507 |
| C | 0.00131 | 0.00387 | 0.00167 | 0.00627 |
| D | 0.00067 | 0.00369 | −0.00018 | 0.00592 |
| E | 0.00042 | 0.00372 | −0.00005 | 0.00541 |
| F | 0.00056 | 0.00358 | −0.00051 | 0.00577 |

(b) Scenario 2: $s^- < 0$ and $s^+ = s^-$.

| | Sel. coeff. $s^-$ | | Sel. coeff. $s^+$ | |
|---|---|---|---|---|
| Scenario | Bias | RMSE | Bias | RMSE |
| A | 0.00503 | 0.00777 | −0.00537 | 0.00896 |
| B | 0.00214 | 0.00585 | −0.00263 | 0.00638 |
| C | 0.00244 | 0.00547 | −0.00331 | 0.00725 |
| D | 0.00128 | 0.00499 | −0.00165 | 0.00593 |
| E | 0.00038 | 0.00462 | −0.00104 | 0.00558 |
| F | 0.00052 | 0.00432 | −0.00018 | 0.00563 |

(c) Scenario 3: $s^- < 0$ and $s^+ > s^-$.

Table S2: Mean bias and RMSE in MAP estimates of the selection coefficients across different data qualities and selection scenarios, corresponding to Figure 3. Mean bias and RMSE are calculated with 200 replicates for each combination. Data qualities (scenarios A–F) are described in Table 1, and selection scenarios (scenarios 1–9) are described in Table 2.

| | Sel. coeff. $s^-$ | | Sel. coeff. $s^+$ | |
|---|---|---|---|---|
| Scenario | Bias | RMSE | Bias | RMSE |
| A | −0.00023 | 0.00347 | 0.00551 | 0.00834 |
| B | 0.00028 | 0.00348 | 0.00191 | 0.00528 |
| C | 0.00022 | 0.00366 | 0.00265 | 0.00571 |
| D | 0.00064 | 0.00362 | −0.00027 | 0.00508 |
| E | −0.00001 | 0.00338 | 0.00079 | 0.00432 |
| F | 0.00063 | 0.00368 | −0.00035 | 0.00424 |

(d) Scenario 4: $s^- = 0$ and $s^+ < s^-$.

| | Sel. coeff. $s^-$ | | Sel. coeff. $s^+$ | |
|---|---|---|---|---|
| Scenario | Bias | RMSE | Bias | RMSE |
| A | 0.00020 | 0.00389 | 0.00026 | 0.00300 |
| B | −0.00001 | 0.00437 | 0.00044 | 0.00337 |
| C | 0.00015 | 0.00368 | 0.00022 | 0.00292 |
| D | 0.00015 | 0.00422 | 0.00040 | 0.00330 |
| E | −0.00003 | 0.00391 | 0.00028 | 0.00301 |
| F | 0.00007 | 0.00380 | 0.00040 | 0.00328 |

(e) Scenario 5: $s^- = 0$ and $s^+ = s^-$.

| | Sel. coeff. $s^-$ | | Sel. coeff. $s^+$ | |
|---|---|---|---|---|
| Scenario | Bias | RMSE | Bias | RMSE |
| A | −0.00031 | 0.00368 | −0.00460 | 0.00722 |
| B | −0.00084 | 0.00409 | −0.00134 | 0.00548 |
| C | −0.00054 | 0.00394 | −0.00195 | 0.00571 |
| D | −0.00061 | 0.00411 | 0.00021 | 0.00548 |
| E | −0.00060 | 0.00365 | −0.00034 | 0.00494 |
| F | −0.00086 | 0.00397 | 0.00037 | 0.00427 |

(f) Scenario 6: $s^- = 0$ and $s^+ > s^-$.

Table S2: Mean bias and RMSE in MAP estimates of the selection coefficients across different data qualities and selection scenarios, corresponding to Figure 3, continued.

| Scenario | Sel. coeff. $s^-$ | | Sel. coeff. $s^+$ | |
|---|---|---|---|---|
| | Bias | RMSE | Bias | RMSE |
| A | −0.00558 | 0.00834 | 0.00585 | 0.00951 |
| B | −0.00259 | 0.00619 | 0.00200 | 0.00611 |
| C | −0.00265 | 0.00580 | 0.00195 | 0.00624 |
| D | −0.00128 | 0.00529 | 0.00021 | 0.00593 |
| E | −0.00064 | 0.00452 | 0.00002 | 0.00569 |
| F | −0.00067 | 0.00475 | 0.00034 | 0.00552 |

(g) Scenario 7: $s^- > 0$ and $s^+ < s^-$.

| Scenario | Sel. coeff. $s^-$ | | Sel. coeff. $s^+$ | |
|---|---|---|---|---|
| | Bias | RMSE | Bias | RMSE |
| A | −0.00295 | 0.00521 | −0.00383 | 0.00721 |
| B | −0.00131 | 0.00415 | −0.00093 | 0.00596 |
| C | −0.00104 | 0.00371 | −0.00219 | 0.00628 |
| D | −0.00066 | 0.00389 | 0.00001 | 0.00547 |
| E | −0.00090 | 0.00385 | −0.00033 | 0.00598 |
| F | −0.00092 | 0.00359 | 0.00079 | 0.00572 |

(h) Scenario 8: $s^- > 0$ and $s^+ = s^-$.

| Scenario | Sel. coeff. $s^-$ | | Sel. coeff. $s^+$ | |
|---|---|---|---|---|
| | Bias | RMSE | Bias | RMSE |
| A | −0.00167 | 0.00412 | −0.00713 | 0.00982 |
| B | −0.00124 | 0.00434 | −0.00294 | 0.00792 |
| C | −0.00142 | 0.00380 | −0.00284 | 0.00744 |
| D | −0.00019 | 0.00350 | −0.00006 | 0.00687 |
| E | −0.00047 | 0.00332 | −0.00082 | 0.00597 |
| F | −0.00038 | 0.00365 | −0.00068 | 0.00612 |

(i) Scenario 9: $s^- > 0$ and $s^+ > s^-$.

Table S2: Mean bias and RMSE in MAP estimates of the selection coefficients across different data qualities and selection scenarios, corresponding to Figure 3, continued.

| Sel. coeff. $s$ | Bias | RMSE |
|---|---|---|
| $s \in [-0.050, -0.010)$ | 0.00106 | 0.00298 |
| $s \in [-0.010, -0.005)$ | 0.00016 | 0.00231 |
| $s \in [-0.005, -0.001)$ | −0.00026 | 0.00205 |
| $s \in [-0.001, 0)$ | 0.00006 | 0.00206 |
| $s \in \{0\}$ | 0.00019 | 0.00201 |
| $s \in (0, 0.001]$ | 0.00020 | 0.00218 |
| $s \in (0.001, 0.005]$ | −0.00003 | 0.00200 |
| $s \in (0.005, 0.010]$ | −0.00015 | 0.00231 |
| $s \in (0.010, 0.050]$ | −0.00077 | 0.00310 |

Table S3: Mean bias and RMSE in MAP estimates of the selection coefficient across different ranges of the selection coefficient $s$ with the parameters $\phi = 0.85$ and $\psi = 1$ (*i.e.*, scenario D in Table 1), corresponding to Figure 4. Mean bias and RMSE are calculated with 200 replicates for each case.

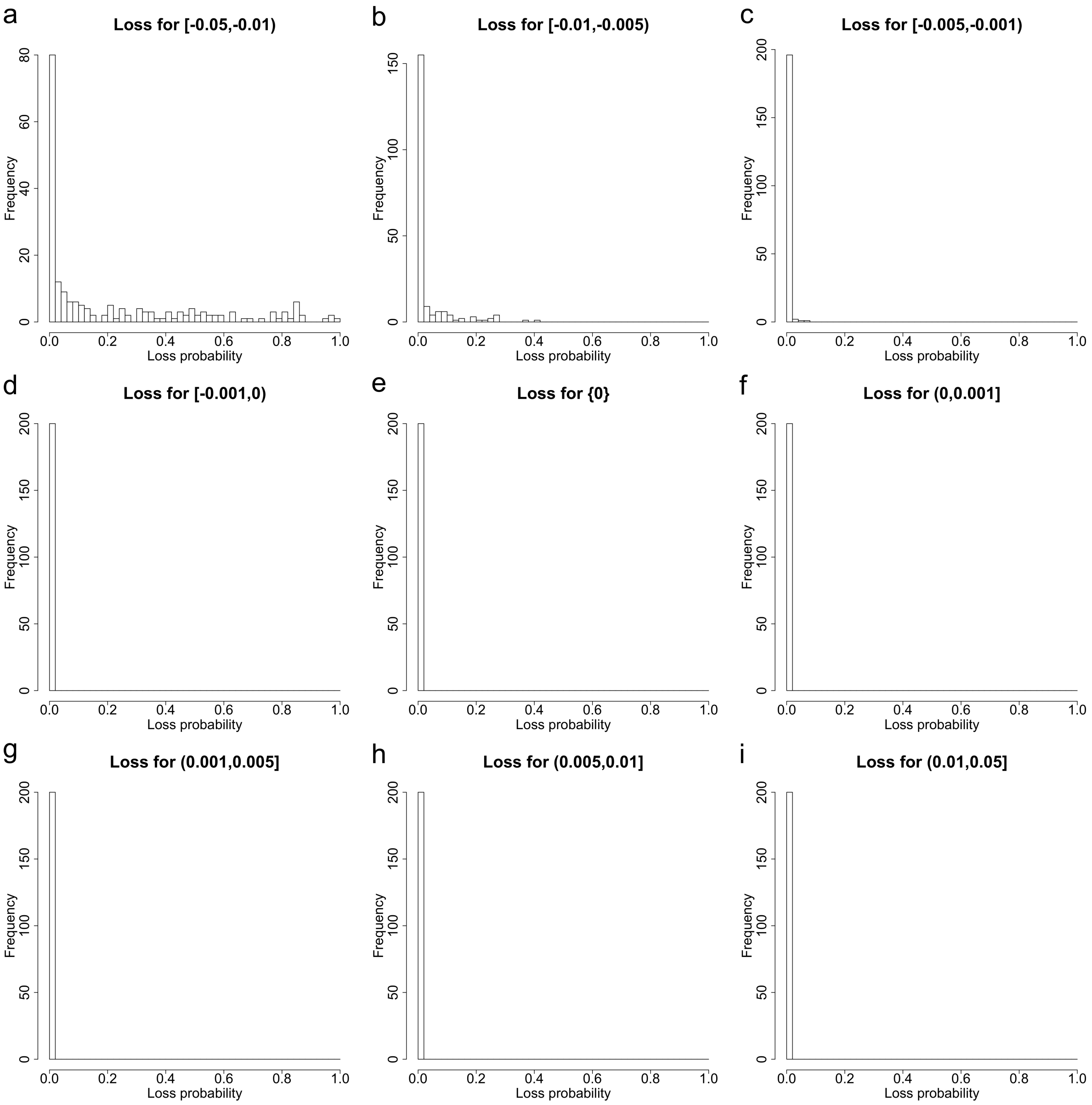


(a) Histograms of loss probabilities.

Figure S1: Empirical distributions for the loss probability and the fixation probability across different ranges of the selection coefficient $s$ with the parameters $\phi = 0.85$ and $\psi = 1$ (*i.e.*, scenario D in Table 1), corresponding to Figure 4.

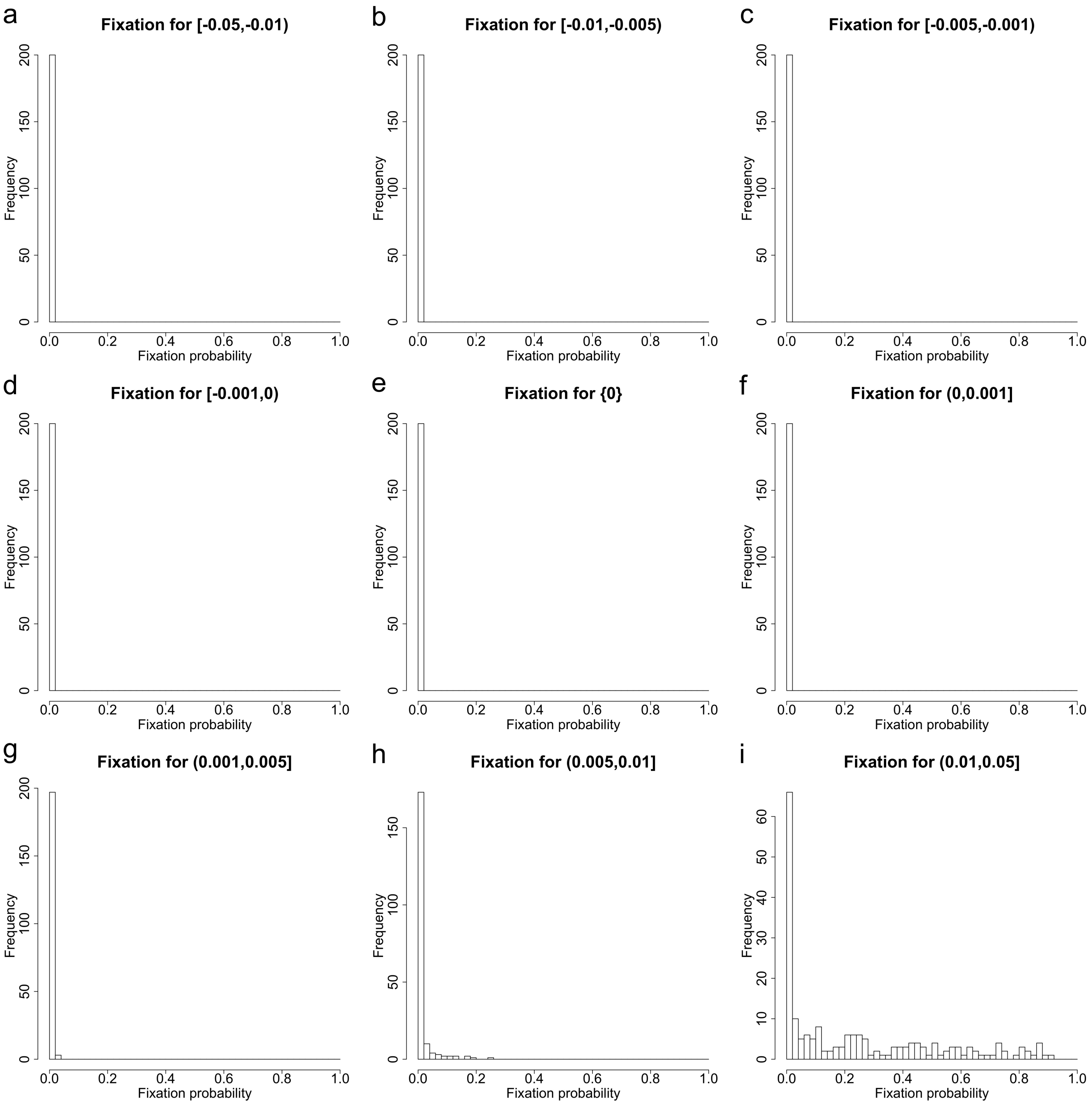


(b) Histograms of fixation probabilities.

Figure S1: Empirical distributions for the loss probability and the fixation probability across different ranges of the selection coefficient $s$ with the parameters $\phi = 0.85$ and $\psi = 1$ (*i.e.*, scenario D in Table 1), corresponding to Figure 4, continued.

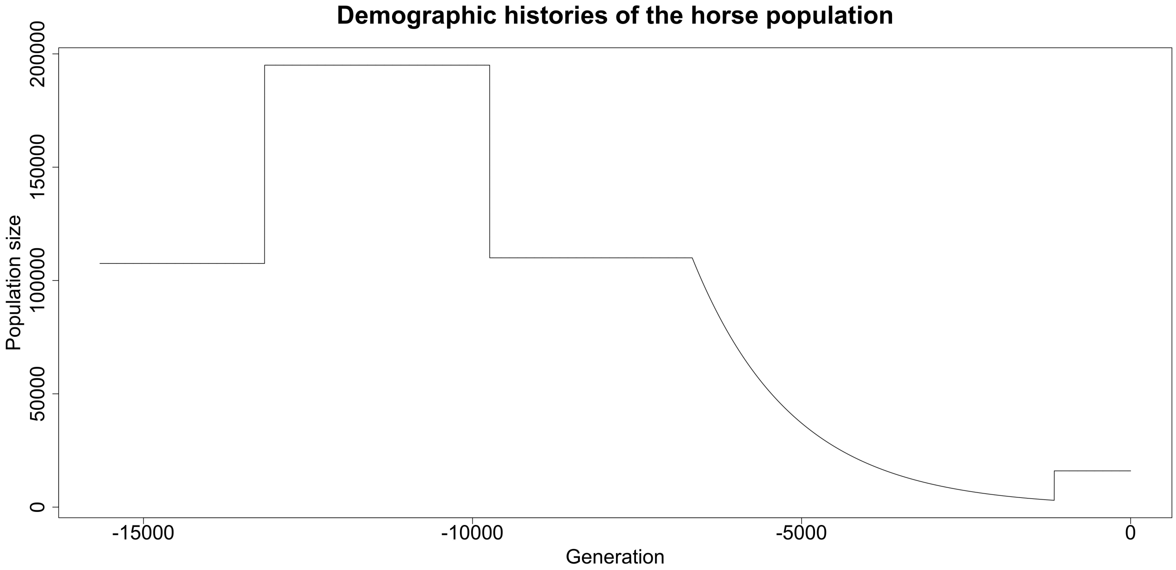


Figure S2: Demographic histories of the horse population reported in Der Sarkissian et al. (2015).

| Gene | Event | Sel. coeff. | MAP est. | 95% HPD | Prob. for -ve. | Prob. for +ve. |
|---|---|---|---|---|---|---|
| | | $s^-$ | 0.00178 | [−0.00120, 0.00660] | 0.0960 | 0.9040 |
| *ASIP* | DOM | $s^+$ | 0.00032 | [−0.00225, 0.00313] | 0.4110 | 0.5890 |
| | | $\Delta s$ | −0.00208 | [−0.00708, 0.00314] | 0.7790 | 0.2210 |
| | | $s^-$ | −0.03003 | [−0.09277, 0.09973] | 0.6695 | 0.3305 |
| *MC1R* | DOM | $s^+$ | 0.01159 | [ 0.00787, 0.01736] | 0.0000 | 1.0000 |
| | | $\Delta s$ | 0.04389 | [−0.08908, 0.10579] | 0.2525 | 0.7475 |
| | | $s^-$ | 0.00392 | [−0.00097, 0.01031] | 0.0650 | 0.9350 |
| *KIT13* | EMA | $s^+$ | −0.02836 | [−0.06270, 0.00152] | 0.9685 | 0.0315 |
| | | $\Delta s$ | −0.03259 | [−0.07003, 0.00078] | 0.9690 | 0.0310 |
| | | $s^-$ | −0.00049 | [−0.00421, 0.00294] | 0.6170 | 0.3830 |
| *TRPM1* | DOM | $s^+$ | −0.00775 | [−0.01387, −0.00046] | 0.9800 | 0.0200 |
| | | $\Delta s$ | −0.00665 | [−0.01419, 0.00250] | 0.9420 | 0.0580 |

Table S5: MAP estimates of the selection coefficients and their changes with their 95% HPD intervals, as well as posterior probabilities for negative selection/change and positive selection/change, for *ASIP*, *MC1R*, *KIT13* and *TRPM1*, corresponding to Figures 7–10. Prob. for -ve. and +ve. stand for posterior probabilities for negative selection/change and positive selection/change, respectively. DOM and EMA stand for domestication and Early Middle Ages, respectively.

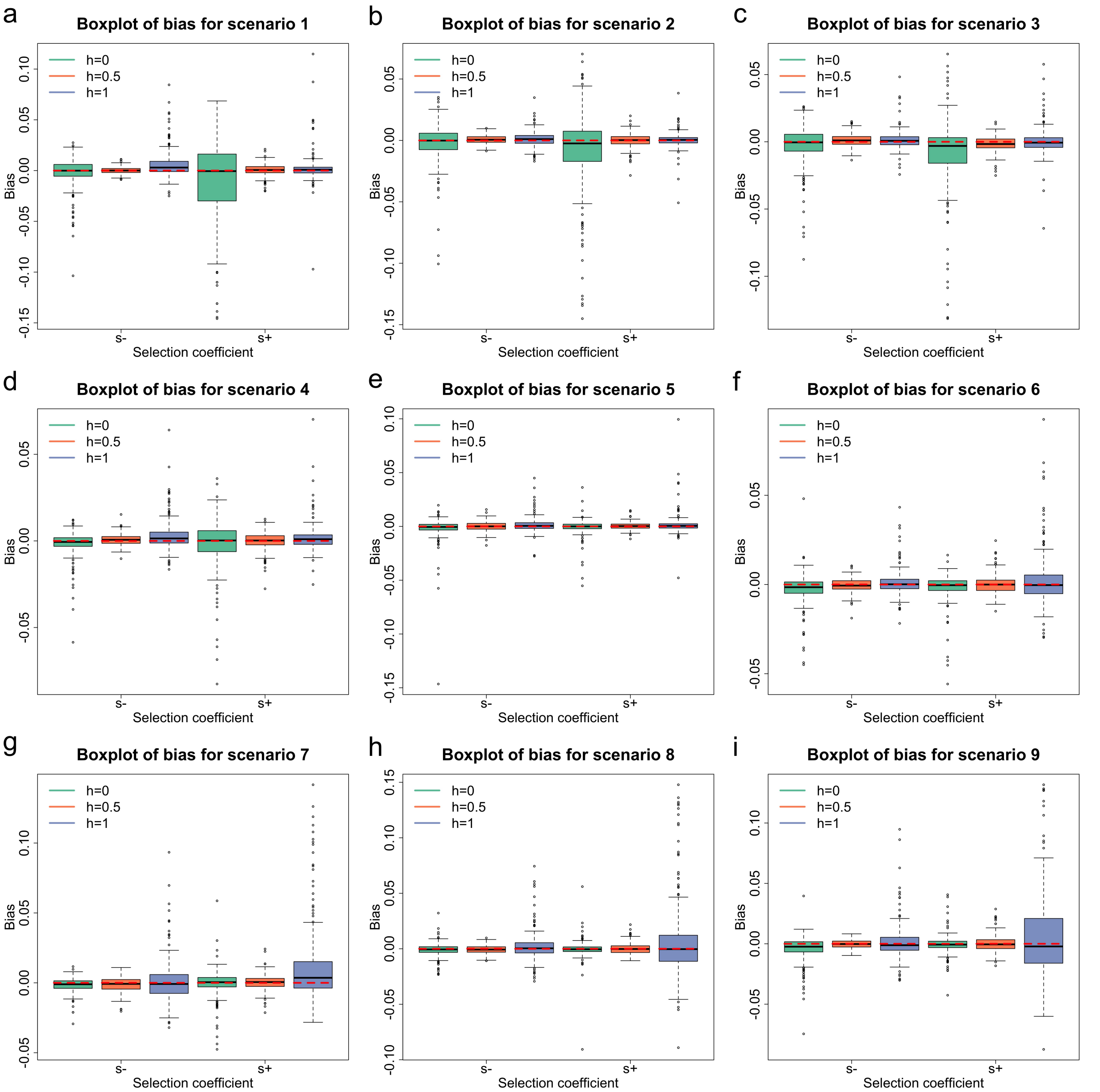


Figure S3: Empirical distributions for the bias in MAP estimates of the selection coefficients across different gene actions and selection scenarios with the parameters $\phi = 0.85$ and $\psi = 1$ (*i.e.*, scenario D in Table 1). For each combination, we repeatedly run our procedure on 200 simulated datasets. Selection scenarios (scenarios 1–9) are described in Table 2. (a)–(i) Boxplots of the bias for scenarios 1–9.

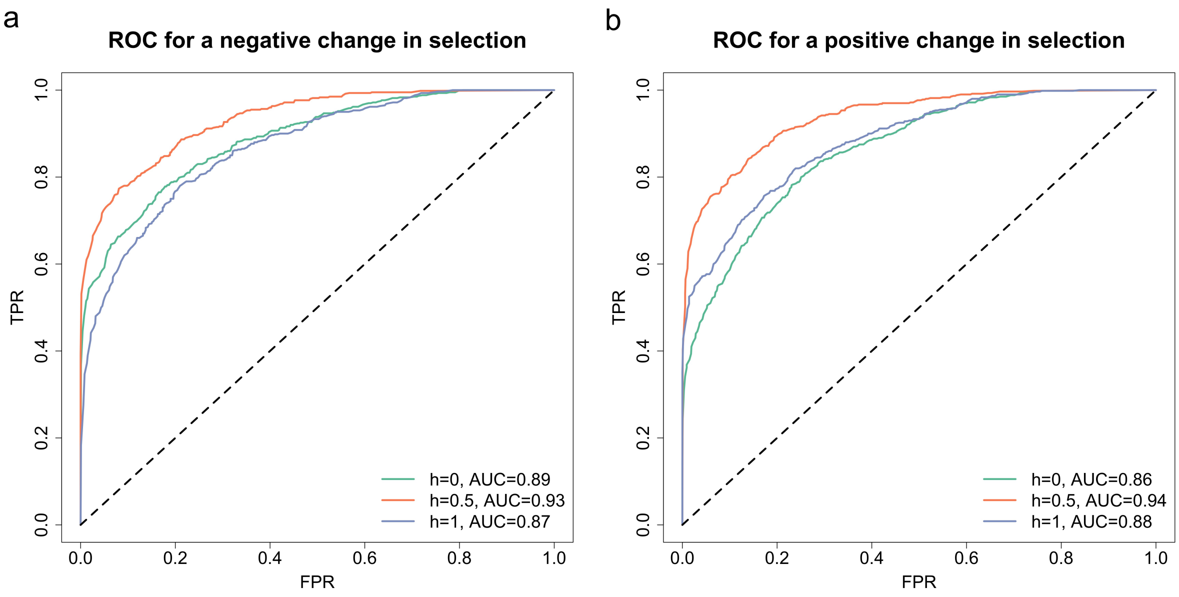


Figure S4: ROC curves for testing a change in selection across different gene actions and selection scenarios with the parameters $\phi = 0.85$ and $\psi = 1$ (*i.e.*, scenario D in Table 1). For each combination, we repeatedly run our procedure on 200 simulated datasets. The AUC value for each curve is summarised. Selection scenarios (scenarios 1–9) are described in Table 2. ROC curves for (a) a negative change in selection and (b) a positive change in selection.

| Dom. param. $h$ | Sel. coeff. $s^-$ | | Sel. coeff. $s^+$ | |
|---|---|---|---|---|
| | Bias | RMSE | Bias | RMSE |
| 0.0 | −0.00254 | 0.01612 | −0.01057 | 0.04167 |
| 0.5 | 0.00022 | 0.00348 | 0.00073 | 0.00595 |
| 1.0 | 0.00614 | 0.01508 | 0.00219 | 0.01537 |

(a) Scenario 1: $s^- < 0$ and $s^+ < s^-$.

| Dom. param. $h$ | Sel. coeff. $s^-$ | | Sel. coeff. $s^+$ | |
|---|---|---|---|---|
| | Bias | RMSE | Bias | RMSE |
| 0.0 | −0.00231 | 0.01663 | −0.00909 | 0.03651 |
| 0.5 | 0.00067 | 0.00369 | −0.00018 | 0.00592 |
| 1.0 | 0.00137 | 0.00633 | 0.00014 | 0.00678 |

(b) Scenario 2: $s^- < 0$ and $s^+ = s^-$.

| Dom. param. $h$ | Sel. coeff. $s^-$ | | Sel. coeff. $s^+$ | |
|---|---|---|---|---|
| | Bias | RMSE | Bias | RMSE |
| 0.0 | −0.00326 | 0.01638 | −0.00951 | 0.03096 |
| 0.5 | 0.00128 | 0.00499 | −0.00165 | 0.00593 |
| 1.0 | 0.00128 | 0.00745 | 0.00026 | 0.01085 |

(c) Scenario 3: $s^- < 0$ and $s^+ > s^-$.

| Dom. param. $h$ | Sel. coeff. $s^-$ | | Sel. coeff. $s^+$ | |
|---|---|---|---|---|
| | Bias | RMSE | Bias | RMSE |
| 0.0 | −0.00181 | 0.00809 | −0.00147 | 0.01473 |
| 0.5 | 0.00064 | 0.00362 | −0.00027 | 0.00508 |
| 1.0 | 0.00334 | 0.00986 | 0.00179 | 0.00866 |

(d) Scenario 4: $s^- = 0$ and $s^+ < s^-$.

| Dom. param. $h$ | Sel. coeff. $s^-$ | | Sel. coeff. $s^+$ | |
|---|---|---|---|---|
| | Bias | RMSE | Bias | RMSE |
| 0.0 | −0.00206 | 0.01300 | −0.00080 | 0.00798 |
| 0.5 | 0.00015 | 0.00422 | 0.00040 | 0.00330 |
| 1.0 | 0.00137 | 0.00727 | 0.00173 | 0.01050 |

(e) Scenario 5: $s^- = 0$ and $s^+ = s^-$.

Table S6: Mean bias and RMSE in MAP estimates of the selection coefficients across different gene actions and selection scenarios with the parameters $\phi = 0.85$ and $\psi = 1$ (*i.e.*, scenario D in Table 1), corresponding to Figure S3. Mean bias and RMSE are calculated with 200 replicates for each combination. Selection scenarios (scenarios 1–9) are described in Table 2.

| Dom. param. $h$ | Sel. coeff. $s^-$ | | Sel. coeff. $s^+$ | |
|---|---|---|---|---|
| | Bias | RMSE | Bias | RMSE |
| 0.0 | −0.00268 | 0.00944 | −0.00176 | 0.00845 |
| 0.5 | −0.00061 | 0.00411 | 0.00021 | 0.00548 |
| 1.0 | 0.00089 | 0.00718 | 0.00222 | 0.01564 |

(f) Scenario 6: $s^- = 0$ and $s^+ > s^-$.

| Dom. param. $h$ | Sel. coeff. $s^-$ | | Sel. coeff. $s^+$ | |
|---|---|---|---|---|
| | Bias | RMSE | Bias | RMSE |
| 0.0 | −0.00139 | 0.00518 | −0.00044 | 0.01025 |
| 0.5 | −0.00128 | 0.00529 | 0.00021 | 0.00593 |
| 1.0 | 0.00120 | 0.01551 | 0.01203 | 0.03144 |

(g) Scenario 7: $s^- > 0$ and $s^+ < s^-$.

| Dom. param. $h$ | Sel. coeff. $s^-$ | | Sel. coeff. $s^+$ | |
|---|---|---|---|---|
| | Bias | RMSE | Bias | RMSE |
| 0.0 | −0.00104 | 0.00632 | −0.00044 | 0.00891 |
| 0.5 | −0.00066 | 0.00389 | 0.00001 | 0.00547 |
| 1.0 | 0.00188 | 0.01367 | 0.00719 | 0.03706 |

(h) Scenario 8: $s^- > 0$ and $s^+ = s^-$.

| Dom. param. $h$ | Sel. coeff. $s^-$ | | Sel. coeff. $s^+$ | |
|---|---|---|---|---|
| | Bias | RMSE | Bias | RMSE |
| 0.0 | −0.00471 | 0.01205 | −0.00043 | 0.00819 |
| 0.5 | −0.00019 | 0.00350 | −0.00006 | 0.00687 |
| 1.0 | 0.00135 | 0.01567 | 0.00558 | 0.03726 |

(i) Scenario 9: $s^- > 0$ and $s^+ > s^-$.

Table S6: Mean bias and RMSE in MAP estimates of the selection coefficients across different gene actions and selection scenarios with the parameters $\phi = 0.85$ and $\psi = 1$ (*i.e.*, scenario D in Table 1), corresponding to Figure S3, continued.

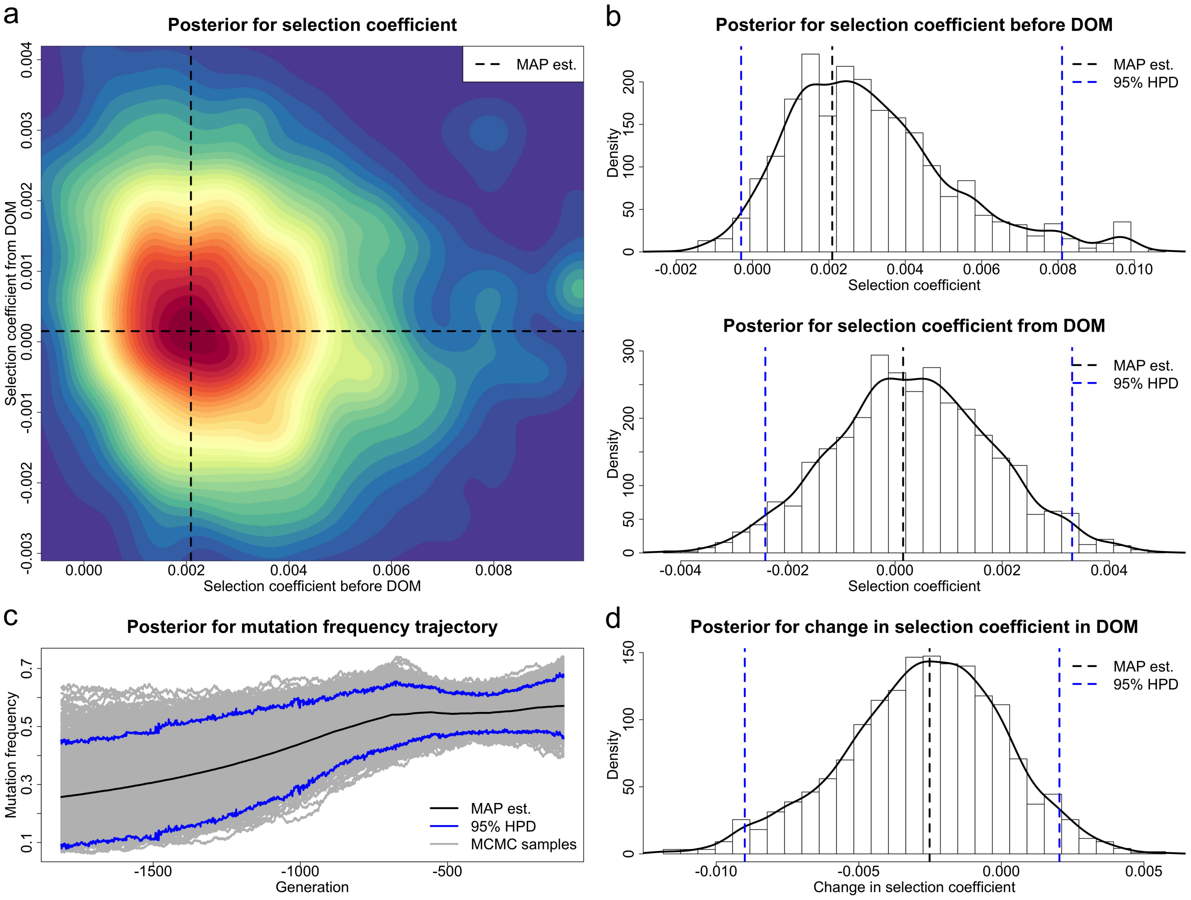


Figure S5: Posteriors for the selection coefficients of the *ASIP* mutation before and from horse domestication (starting from 3500 BC) and the underlying frequency trajectory of the *ASIP* mutation in the population. The demographic history is fixed at $N = 16000$, and the samples drawn before 12500 BC are excluded. DOM stands for domestication. (a) Joint posterior for the selection coefficients $s^-$ and $s^+$. (b) Marginal posteriors for the selection coefficients $s^-$ and $s^+$. (c) Posterior for the underlying frequency trajectory of the *ASIP* mutation. (d) Posterior for the selection change $\Delta s$.

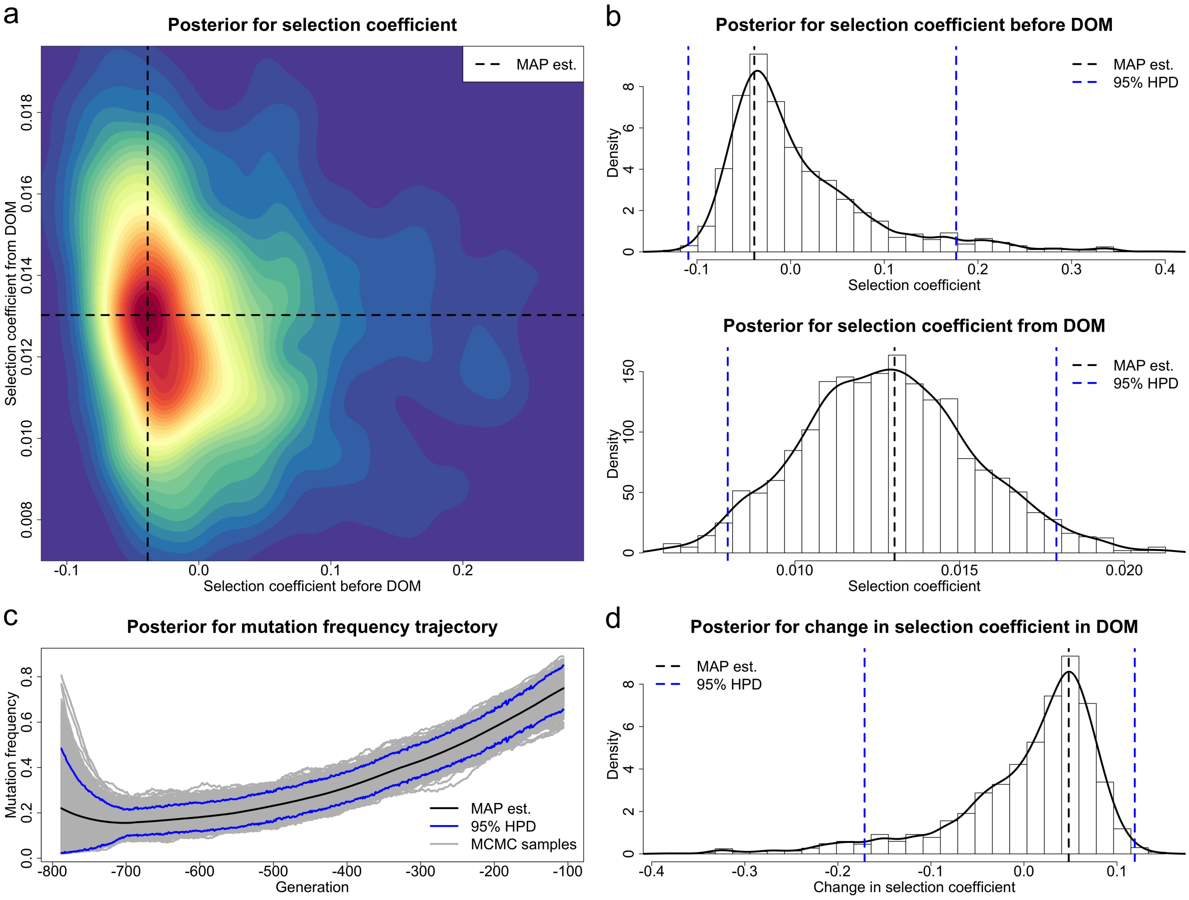


Figure S6: Posteriors for the selection coefficients of the *MC1R* mutation before and from horse domestication (starting from 3500 BC) and the underlying frequency trajectory of the *MC1R* mutation in the population. The demographic history is fixed at $N = 16000$, and the samples drawn before 4300 BC are excluded. DOM stands for domestication. (a) Joint posterior for the selection coefficients $s^-$ and $s^+$. (b) Marginal posteriors for the selection coefficients $s^-$ and $s^+$. (c) Posterior for the underlying frequency trajectory of the *MC1R* mutation. (d) Posterior for the selection change $\Delta s$.

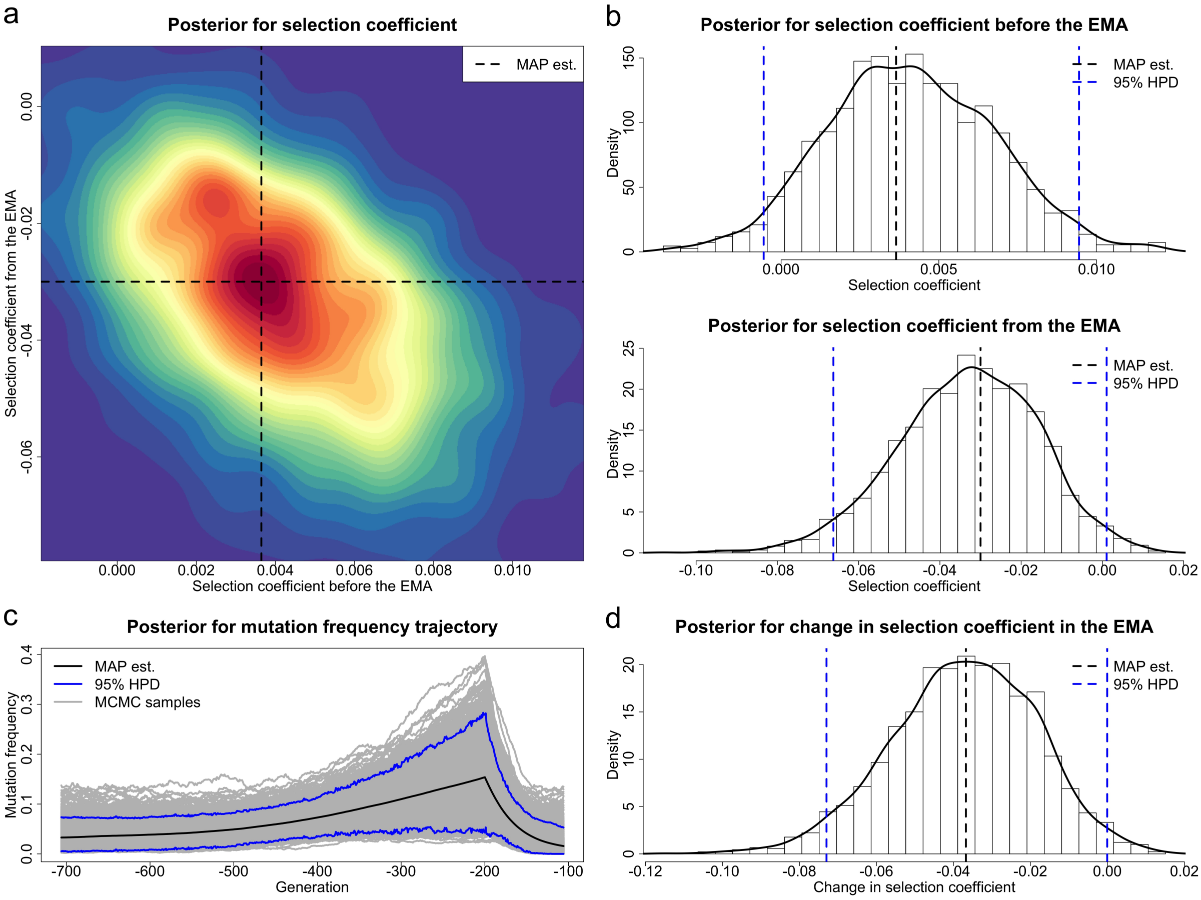


Figure S7: Posteriors for the selection coefficients of the *KIT13* mutation before and from the Middle Ages (starting from AD 400) and the underlying frequency trajectory of the *KIT13* mutation in the population. The demographic history is fixed at $N = 16000$, and the samples drawn before 3645 BC are excluded. EMA stands for Early Middle Ages. (a) Joint posterior for the selection coefficients $s^-$ and $s^+$. (b) Marginal posteriors for the selection coefficients $s^-$ and $s^+$. (c) Posterior for the underlying frequency trajectory of the *KIT13* mutation. (d) Posterior for the selection change $\Delta s$.

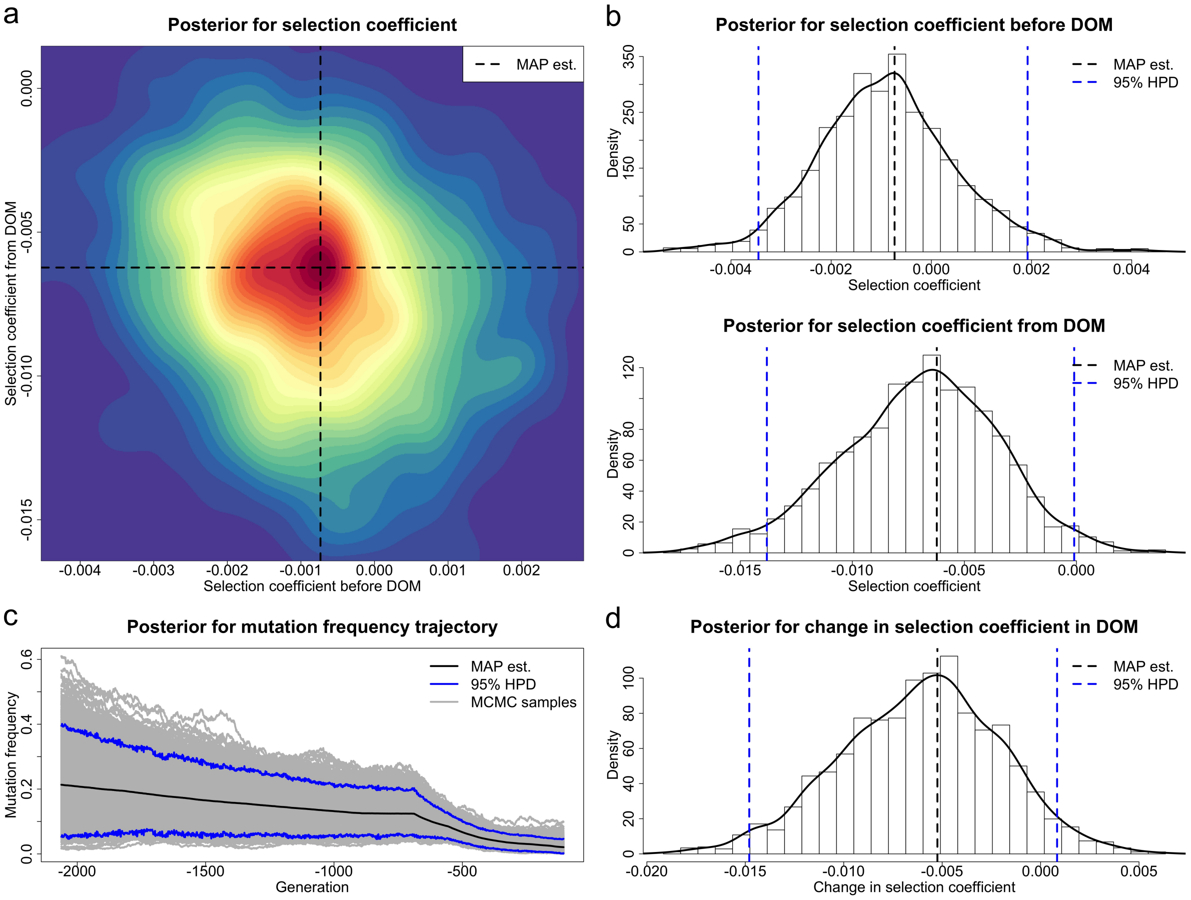


Figure S8: Posteriors for the selection coefficients of the *TRPM1* mutation before and from horse domestication (starting from 3500 BC) and the underlying frequency trajectory of the *TRPM1* mutation in the population. The demographic history is fixed at $N = 16000$, and the samples drawn before 14500 BC are excluded. DOM stands for domestication. (a) Joint posterior for the selection coefficients $s^-$ and $s^+$. (b) Marginal posteriors for the selection coefficients $s^-$ and $s^+$. (c) Posterior for the underlying frequency trajectory of the *TRPM1* mutation. (d) Posterior for the selection change $\Delta s$.

| Gene | Event | Sel. coeff. | MAP est. | 95% HPD | Prob. for -ve. | Prob. for +ve. |
|---|---|---|---|---|---|---|
| | | $s^-$ | 0.00209 | [−0.00030, 0.00810] | 0.0370 | 0.9630 |
| *ASIP* | DOM | $s^+$ | 0.00015 | [−0.00242, 0.00331] | 0.4195 | 0.5805 |
| | | $\Delta s$ | −0.00252 | [−0.00900, 0.00204] | 0.8455 | 0.1545 |
| | | $s^-$ | −0.03878 | [−0.10923, 0.17673] | 0.5920 | 0.4080 |
| *MC1R* | DOM | $s^+$ | 0.01303 | [ 0.00796, 0.01794] | 0.0000 | 1.0000 |
| | | $\Delta s$ | 0.04813 | [−0.17128, 0.11910] | 0.3510 | 0.6490 |
| | | $s^-$ | 0.00364 | [−0.00055, 0.00944] | 0.0525 | 0.9475 |
| *KIT13* | EMA | $s^+$ | −0.03000 | [−0.06621, 0.00104] | 0.9805 | 0.0195 |
| | | $\Delta s$ | −0.03676 | [−0.07297, −0.00005] | 0.9840 | 0.0160 |
| | | $s^-$ | −0.00073 | [−0.00345, 0.00192] | 0.7635 | 0.2365 |
| *TRPM1* | DOM | $s^+$ | −0.00623 | [−0.01381, −0.00010] | 0.9830 | 0.0170 |
| | | $\Delta s$ | −0.00524 | [−0.01480, 0.00084] | 0.9440 | 0.0560 |

Table S7: MAP estimates of the selection coefficients and their changes with their 95% HPD intervals, as well as posterior probabilities for negative selection/change and positive selection/change, for *ASIP*, *MC1R*, *KIT13* and *TRPM1*, corresponding to Figures S5–S8. Prob. for -ve. and +ve. stand for posterior probabilities for negative selection/change and positive selection/change, respectively. The demographic history is fixed at $N = 16000$. DOM and EMA stand for domestication and Early Middle Ages, respectively.

| Method | Data | Gene | $s$ | $h$ | $N$ | Reject $s = 0$ | Note |
|---|---|---|---|---|---|---|---|
| Bollback et al. (2008) | Ludwig et al. (2009) | *ASIP* | $s > 0$ | codominance | constant | yes | 5 years/generation, 6 sampling times |
| | | *MC1R* | $s > 0$ | codominance | constant | yes | 5 years/generation, 6 sampling times |
| Malaspinas et al. (2012) | Ludwig et al. (2009) | *ASIP* | $s < 0$ | recessive | constant | no | 5 years/generation, 6 sampling times |
| Steinrücken et al. (2014) | Ludwig et al. (2009) | *ASIP* | $s > 0$ | overdominance | constant | yes | 5 years/generation, 6 sampling times |
| | | *MC1R* | $s > 0$ | overdominance | constant | yes | 5 years/generation, 6 sampling times |
| Schraiber et al. (2016) | Ludwig et al. (2009) | *ASIP* | $s > 0$ | codominance | time-varying | yes | 8 years/generation, 6 sampling times |
| | | *MC1R* | $s > 0$ | overdominance | time-varying | yes | 8 years/generation, 6 sampling times |
| He et al. (2020a) | Wutke et al. (2018) | *KIT13* | $s > 0$ | dominant | constant | no | 8 years/generation, 9 sampling times |
| He et al. (2020b) | Wutke et al. (2018) | *ASIP* | $s > 0$ | recessive | time-varying | yes | 8 years/generation, 65 sampling times |
| | | *MC1R* | $s > 0$ | recessive | time-varying | yes | 8 years/generation, 65 sampling times |

Table S8: Summary of the results for selection acting on the horse coat colouration loci produced from ancient horse samples in earlier studies. Among these studies, except for Steinrücken et al. (2014) and Schraiber et al. (2016), the dominance parameter $h$ was prespecified, and except for Malaspinas et al. (2012), the population size $N$ was prespecified. The different constant population size was adopted across different studies, and the same time-varying population size reported in Der Sarkissian et al. (2015) was used in Schraiber et al. (2016) and He et al. (2020b). The selection coefficient of *KIT13* in He et al. (2020a) was co-estimated with that of *KIT16* in consideration of genetic recombination and local linkage. For comparative purposes, the estimates for the selection coefficients are all converted into the same selection model parameterisation adopted in our work. The selection coefficient $s$ represents the strength of selection acting on the homozygous double mutant genotype except for $ASIP$ in Steinrücken et al. (2014), where $s$ denotes the strength of selection acting on the heterozygous genotype since their estimate for the homozygous double mutant genotype is 0. Note that the dataset of Wutke et al. (2018) consists of the ancient horse samples from Ludwig et al. (2009), Pruvost et al. (2011) and Wutke et al. (2018).

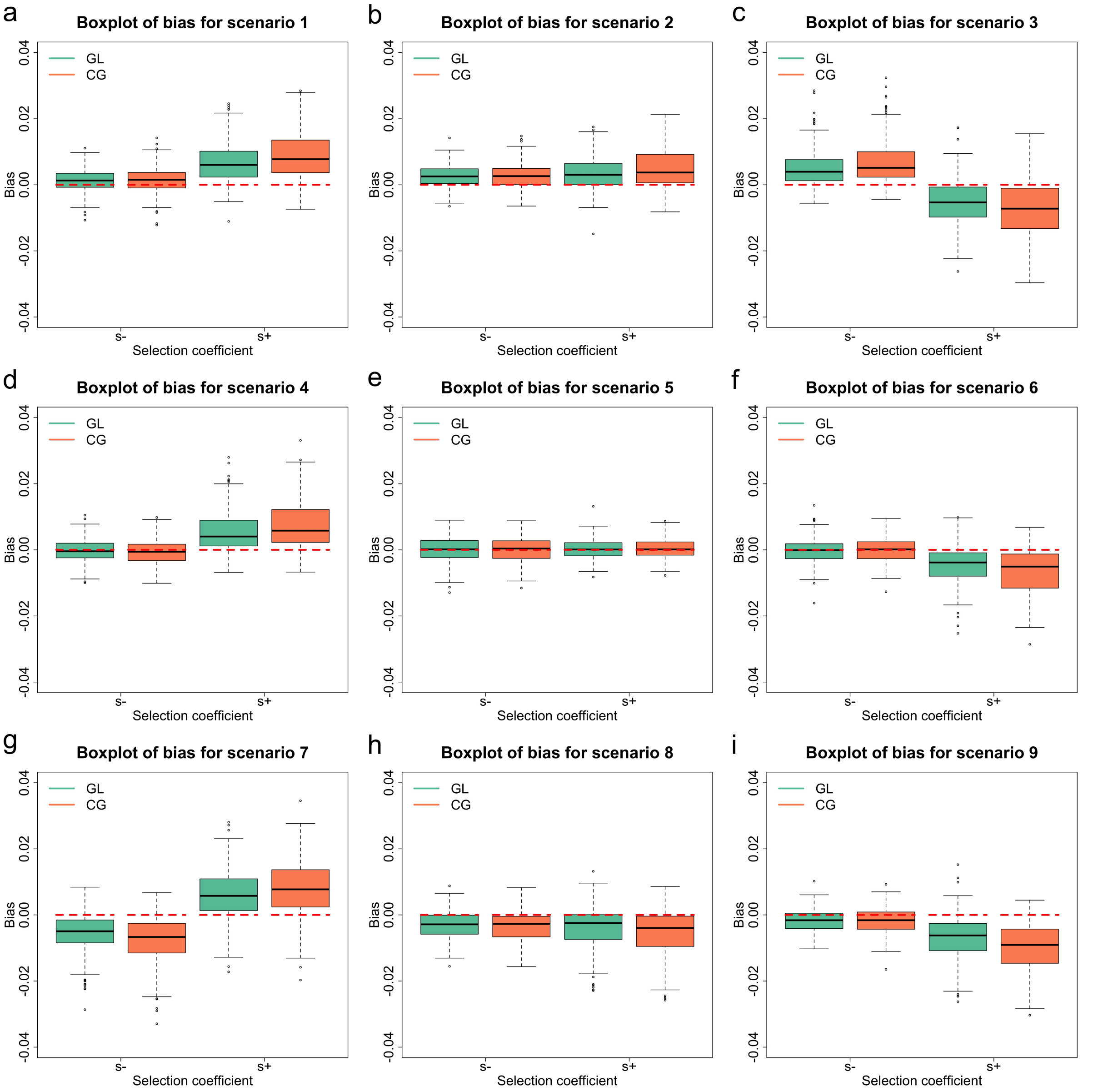


(a) Scenario A: $\phi = 0.75$ and $\psi = 0.5$.

Figure S9: Empirical distributions for the bias in MAP estimates of the selection coefficients across different input types, data qualities and selection scenarios. For each combination, we repeatedly run our procedure on 200 simulated datasets. Data qualities (scenarios A–F) are described in Table 1, and selection scenarios (scenarios 1–9) are described in Table 2. GL and CG are shorthands separately for genotype likelihood and called genotype.

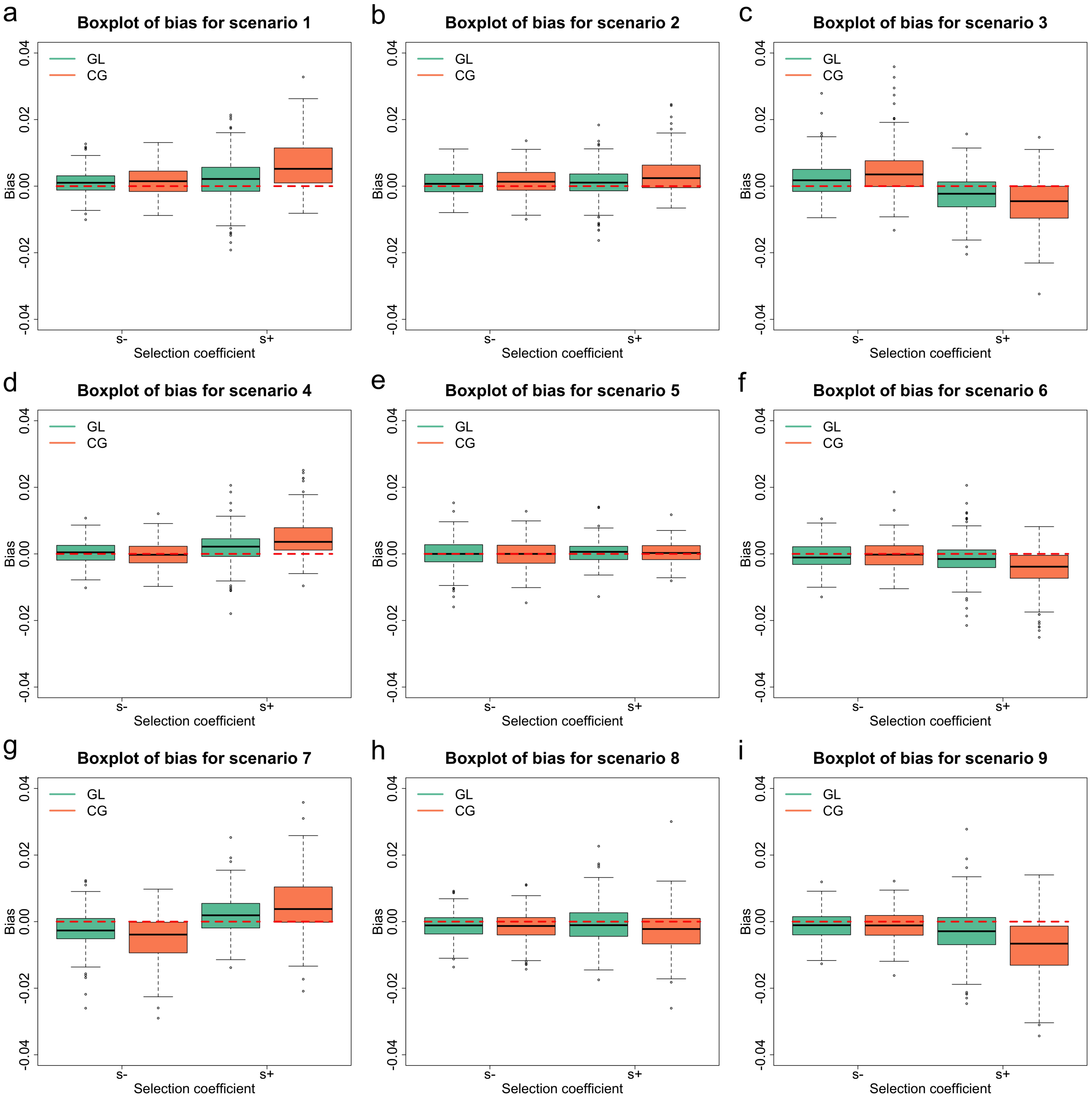


(b) Scenario B: $\phi = 0.75$ and $\psi = 1$.

Figure S9: Empirical distributions for the bias in MAP estimates of the selection coefficients across different input types, data qualities and selection scenarios, continued.

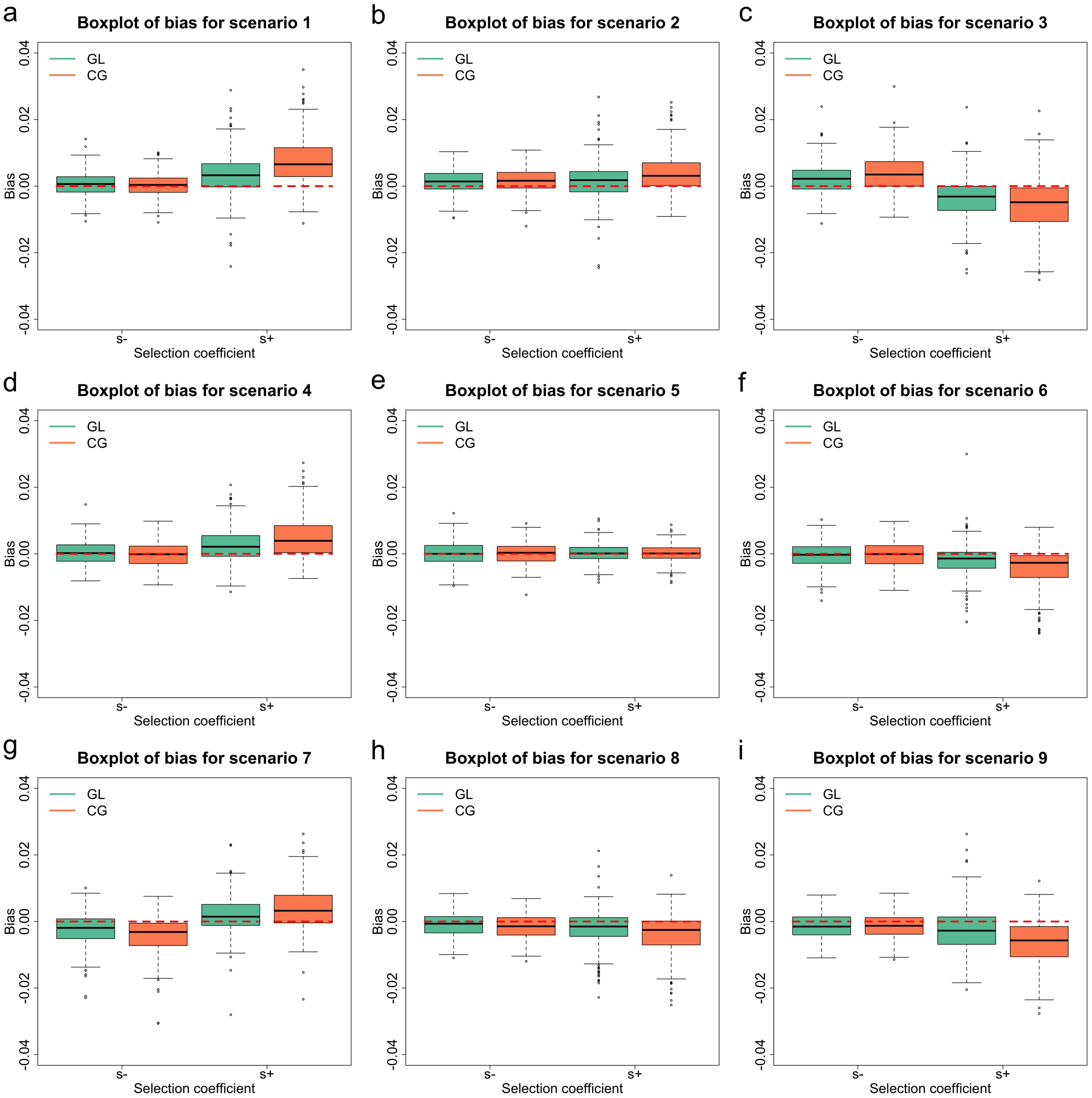


(c) Scenario C: $\phi = 0.85$ and $\psi = 0.5$.

Figure S9: Empirical distributions for the bias in MAP estimates of the selection coefficients across different input types, data qualities and selection scenarios, continued.

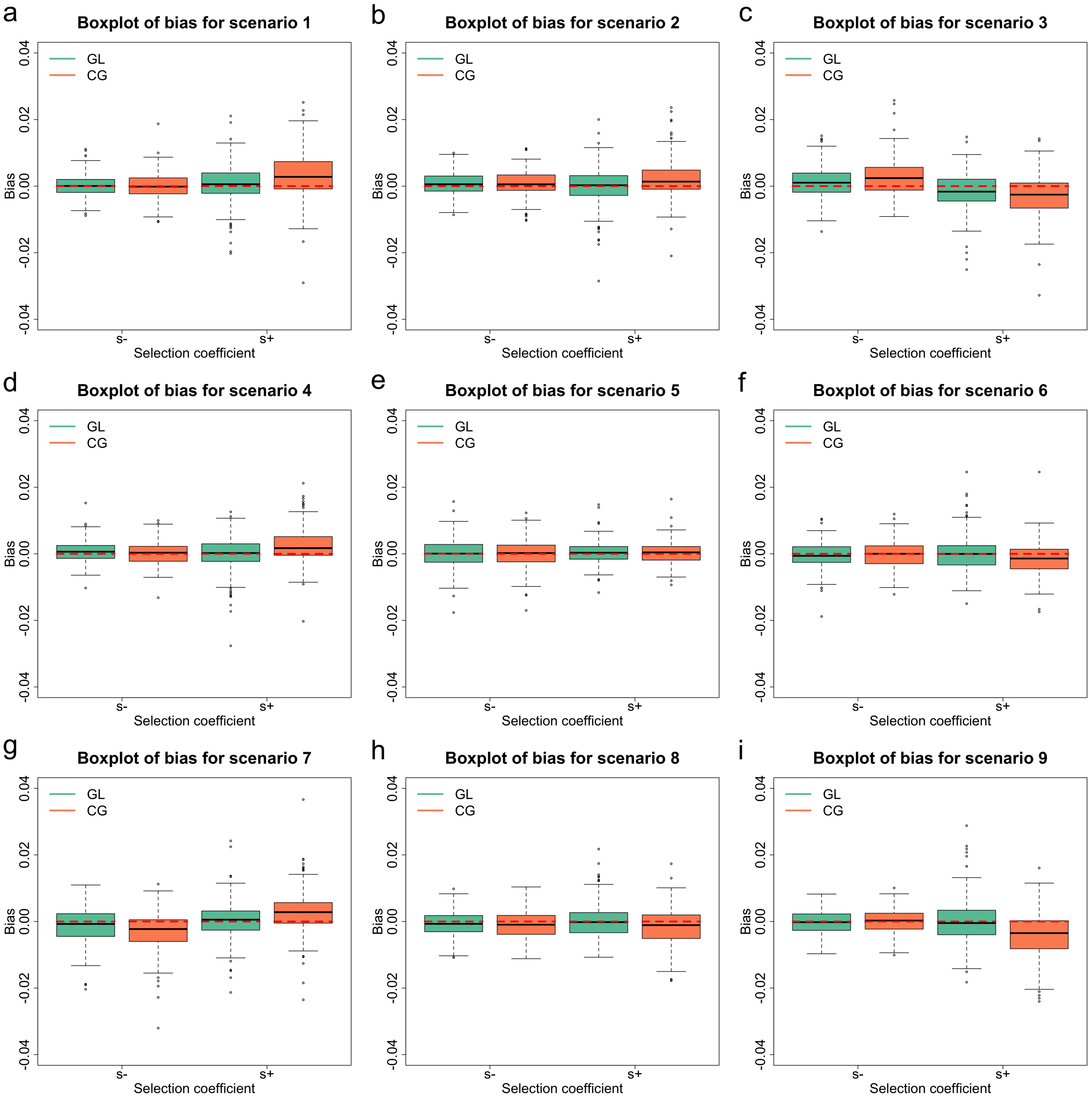


(d) Scenario D: $\phi = 0.85$ and $\psi = 1$.

Figure S9: Empirical distributions for the bias in MAP estimates of the selection coefficients across different input types, data qualities and selection scenarios, continued.

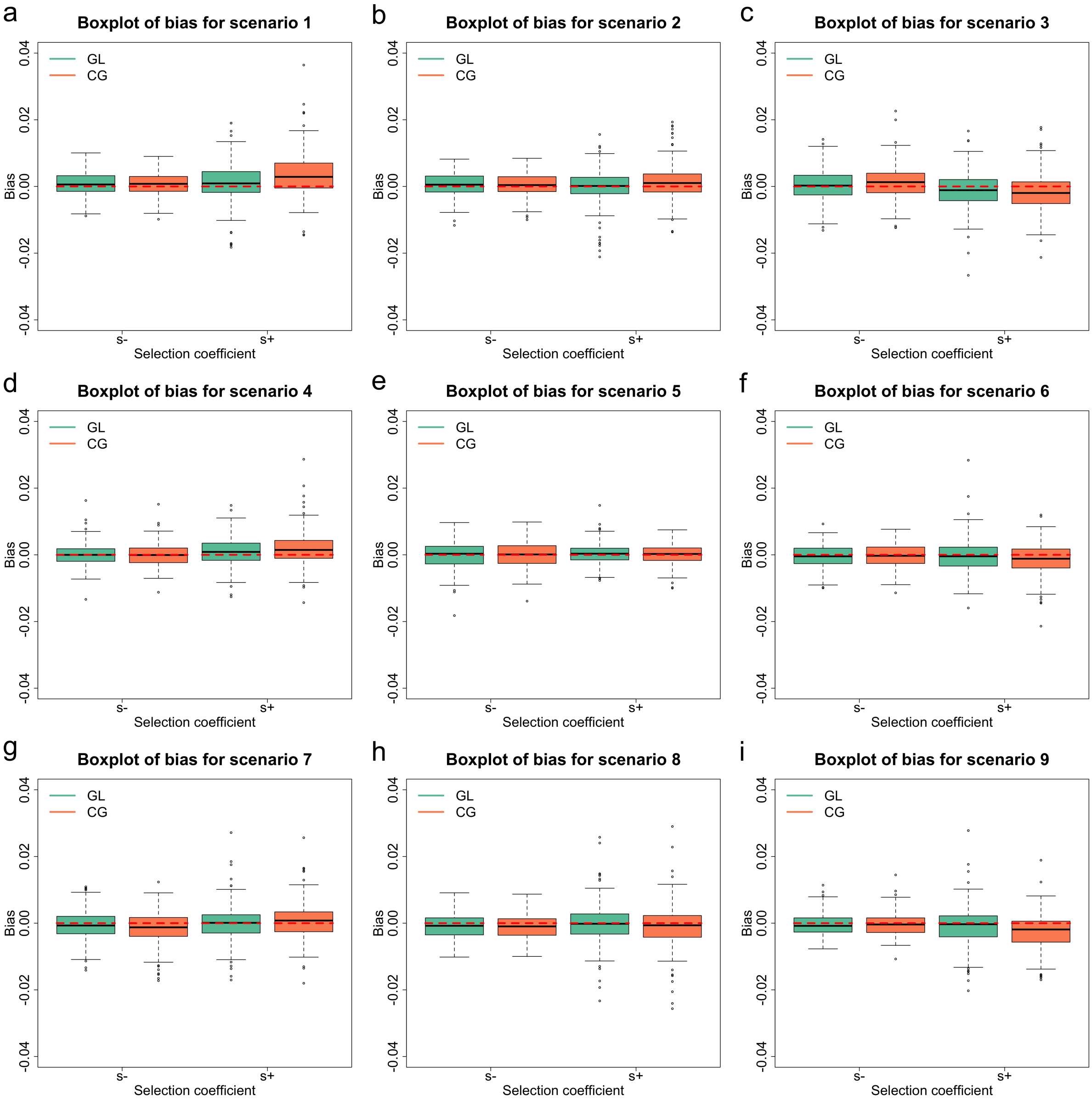


(e) Scenario E: $\phi = 0.95$ and $\psi = 0.5$.

Figure S9: Empirical distributions for the bias in MAP estimates of the selection coefficients across different input types, data qualities and selection scenarios, continued.

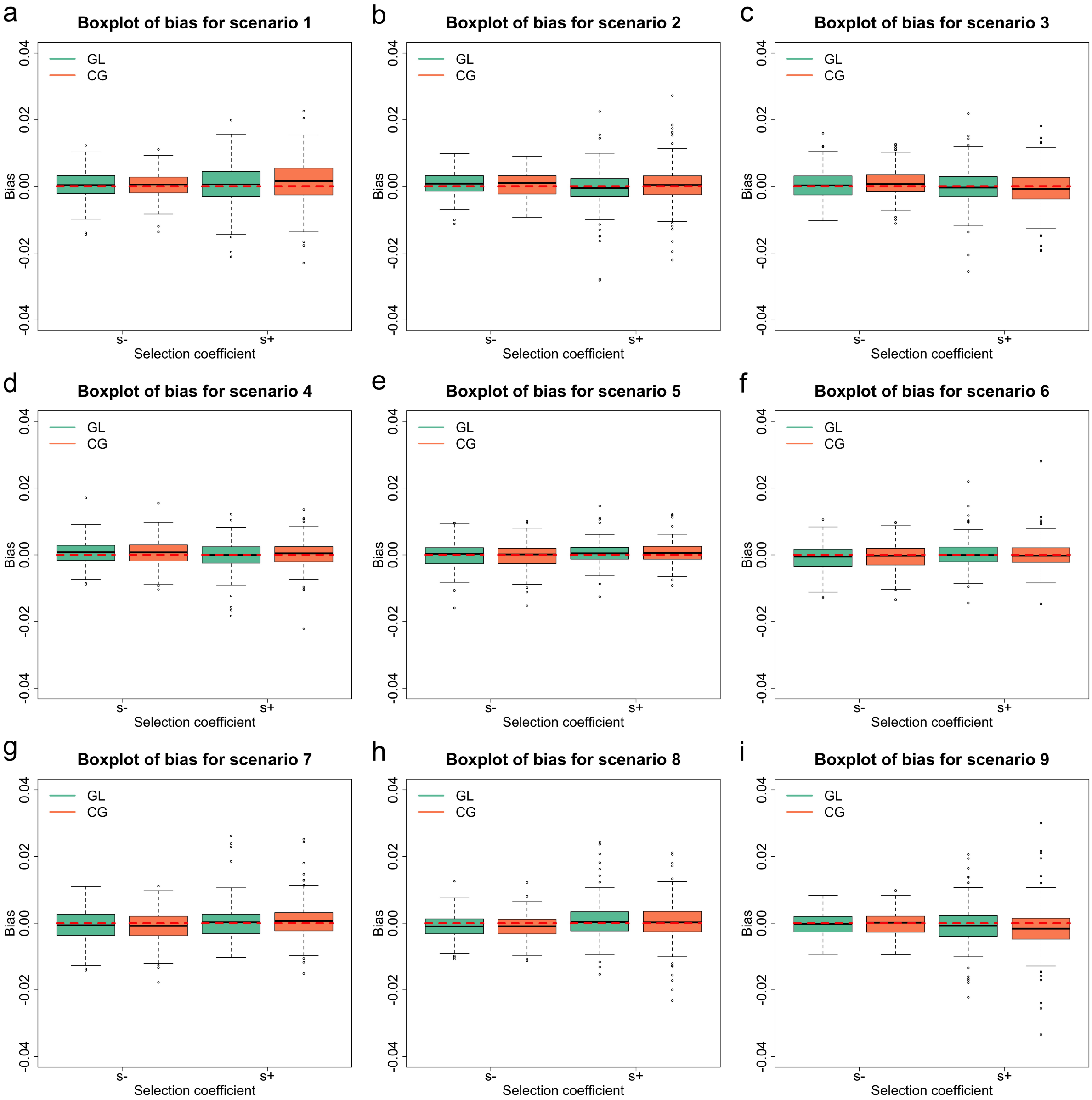


(f) Scenario F: $\phi = 0.95$ and $\psi = 1$.

Figure S9: Empirical distributions for the bias in MAP estimates of the selection coefficients across different input types, data qualities and selection scenarios, continued.

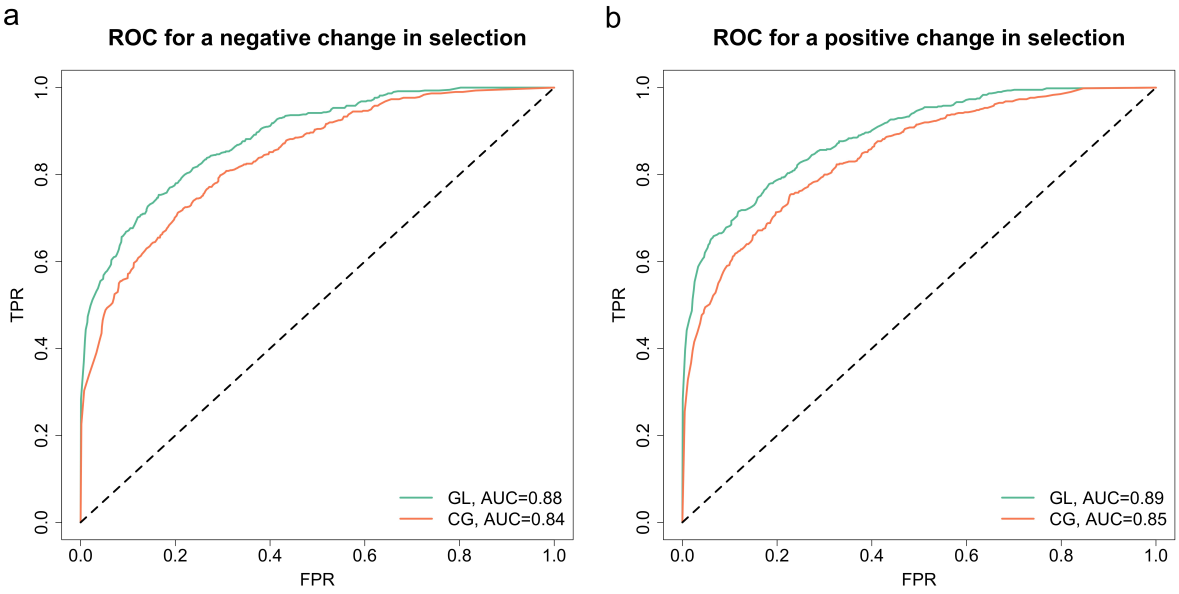


(a) Scenario A: $\phi = 0.75$ and $\psi = 0.5$.

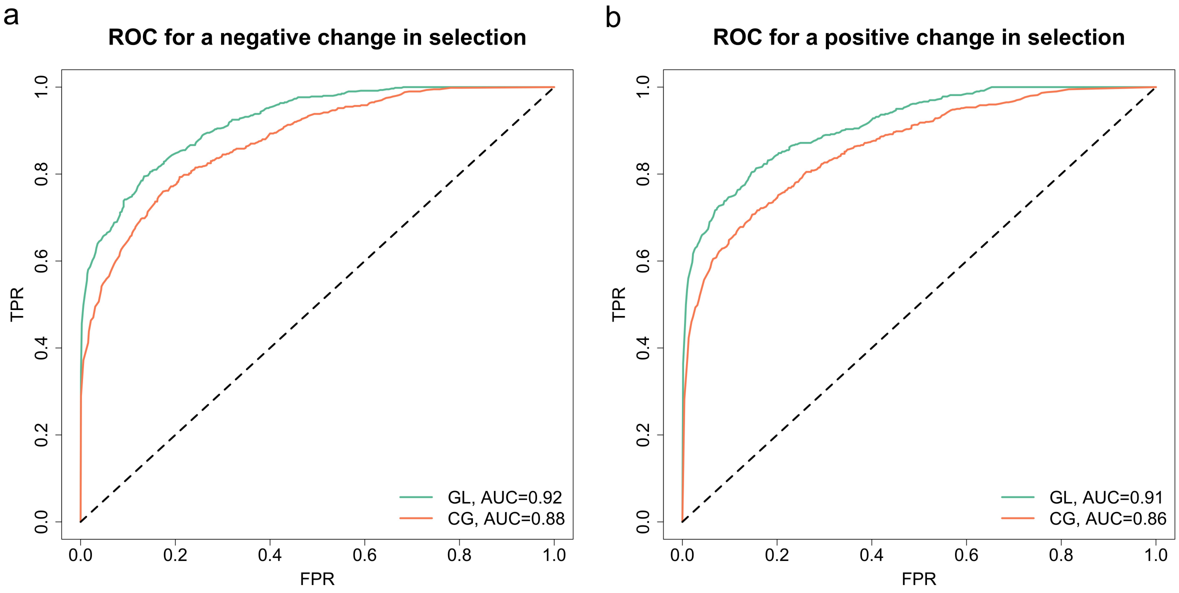


(b) Scenario B: $\phi = 0.75$ and $\psi = 1$.

Figure S10: ROC curves for testing a change in selection across different input types, data qualities and selection scenarios. For each combination, we repeatedly run our procedure on 200 simulated datasets. The AUC value for each curve is summarised. Data qualities (scenarios A–F) are described in Table 1, and selection scenarios (scenarios 1–9) are described in Table 2. GL and CG are shorthands separately for genotype likelihood and called genotype.

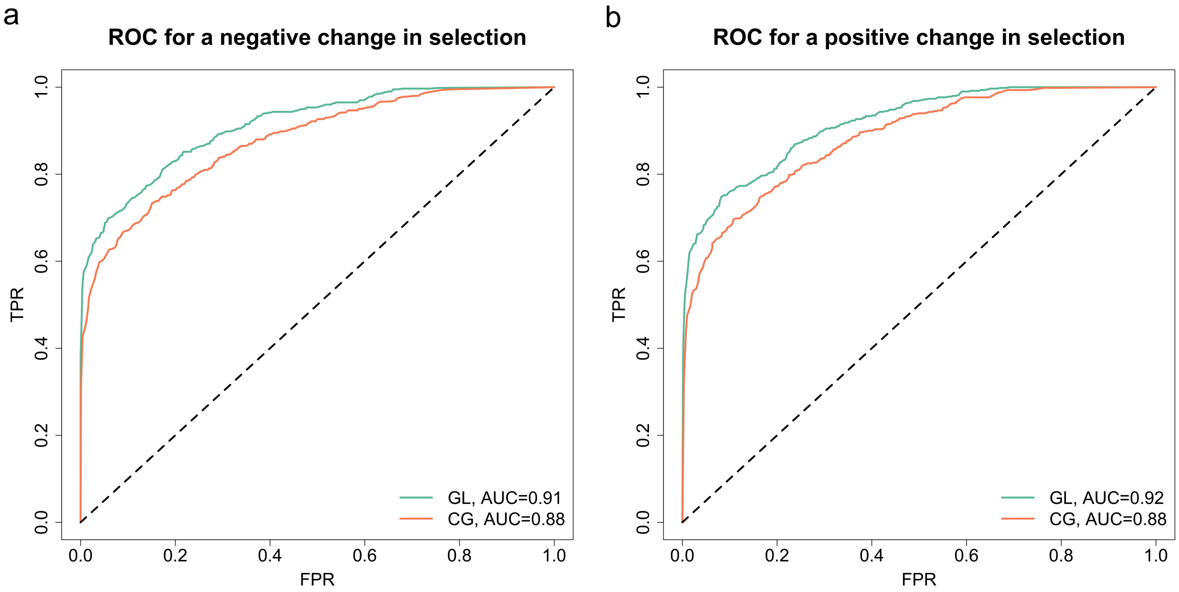


(c) Scenario C: $\phi = 0.85$ and $\psi = 0.5$.

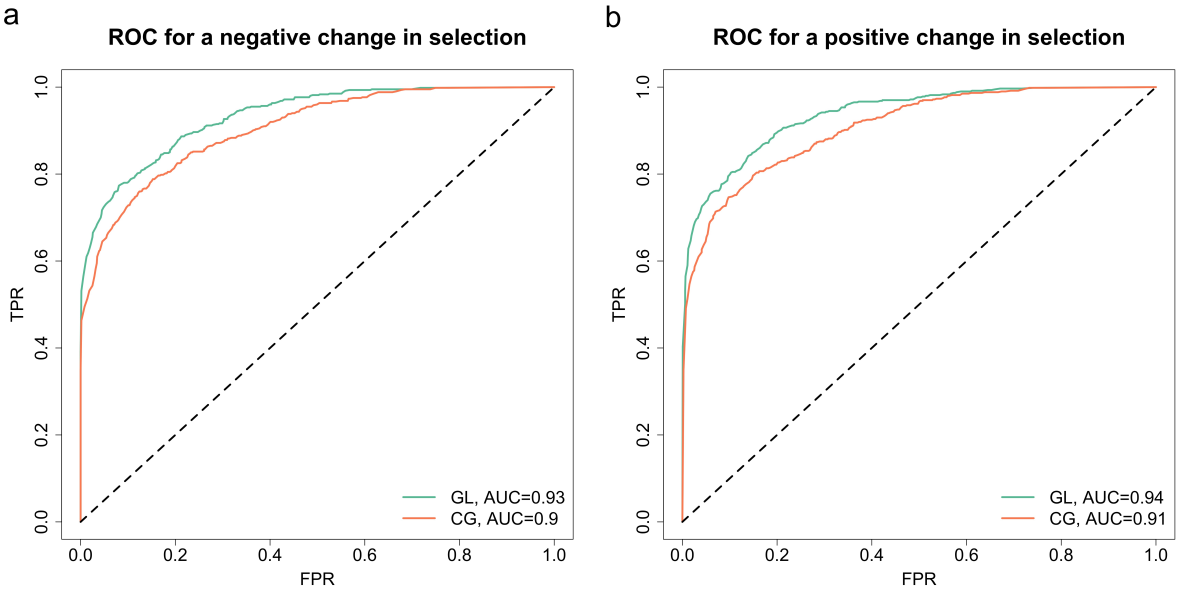


(d) Scenario D: $\phi = 0.85$ and $\psi = 1$.

Figure S10: ROC curves for testing a change in selection across different input types, data qualities and selection scenarios, continued.

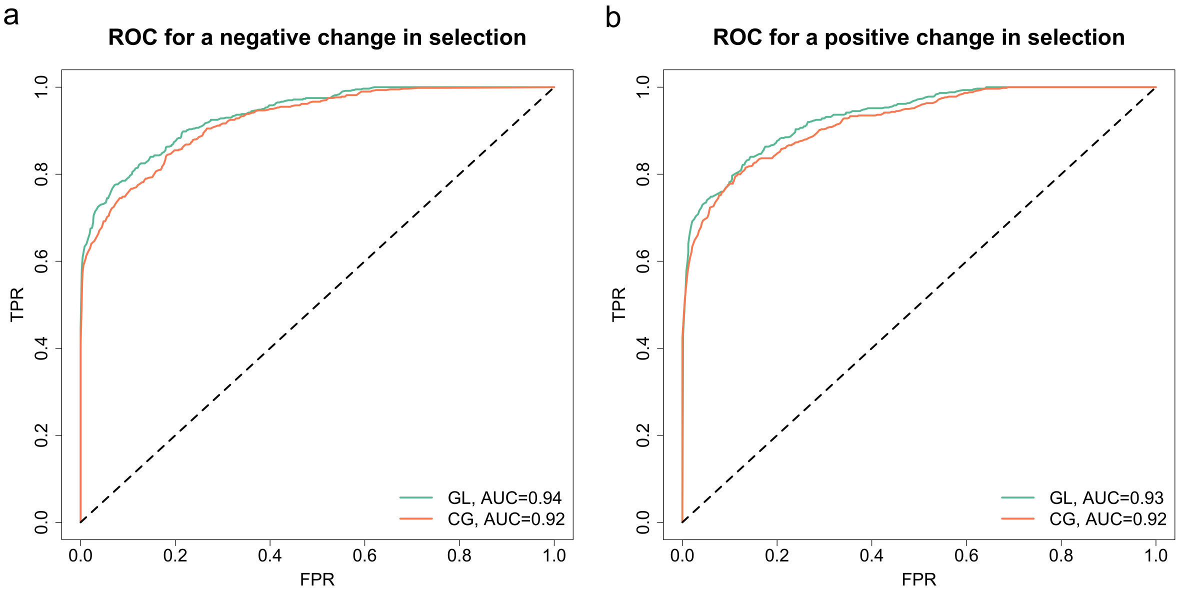


(e) Scenario E: $\phi = 0.95$ and $\psi = 0.5$.

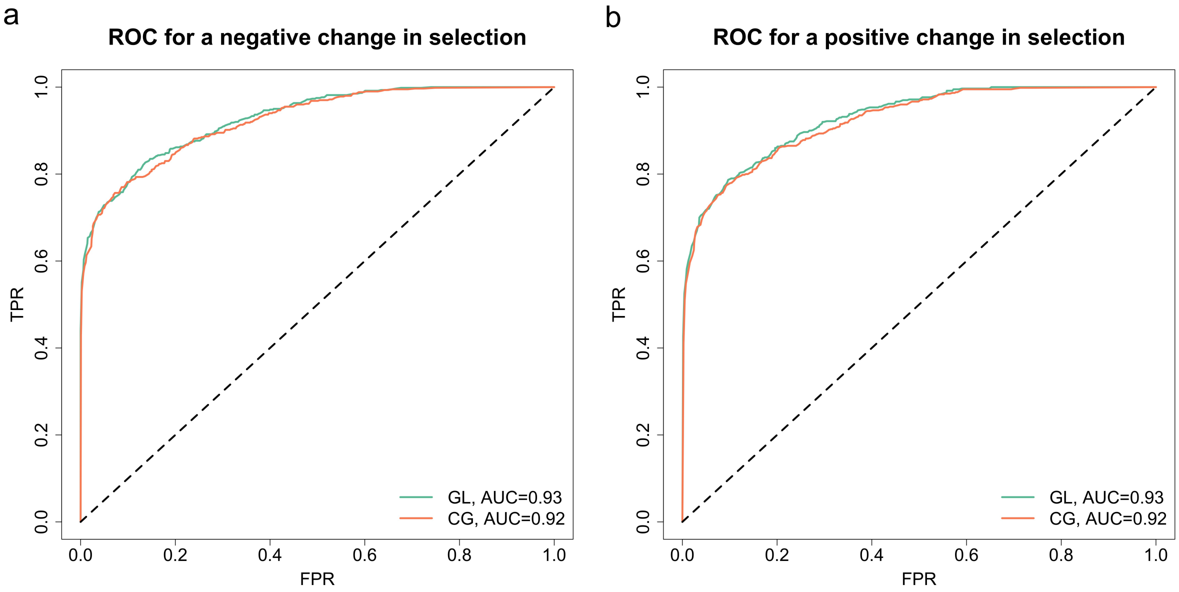


(f) Scenario F: $\phi = 0.95$ and $\psi = 1$.

Figure S10: ROC curves for testing a change in selection across different input types, data qualities and selection scenarios, continued.

| Input | Scenario | Sel. coeff. $s^-$ Bias | Sel. coeff. $s^-$ RMSE | Sel. coeff. $s^+$ Bias | Sel. coeff. $s^+$ RMSE |
|---|---|---|---|---|---|
| GL | A | 0.00126 | 0.00391 | 0.00684 | 0.00952 |
| | B | 0.00124 | 0.00464 | 0.00229 | 0.00708 |
| | C | 0.00054 | 0.00399 | 0.00378 | 0.00806 |
| | D | 0.00022 | 0.00348 | 0.00073 | 0.00595 |
| | E | 0.00081 | 0.00354 | 0.00104 | 0.00603 |
| | F | 0.00043 | 0.00402 | 0.00053 | 0.00618 |
| CG | A | 0.00142 | 0.00427 | 0.00920 | 0.01185 |
| | B | 0.00157 | 0.00467 | 0.00671 | 0.01028 |
| | C | 0.00037 | 0.00380 | 0.00779 | 0.01082 |
| | D | 0.00008 | 0.00384 | 0.00357 | 0.00754 |
| | E | 0.00061 | 0.00358 | 0.00364 | 0.00753 |
| | F | 0.00038 | 0.00374 | 0.00153 | 0.00659 |

(a) Scenario 1: $s^- < 0$ and $s^+ < s^-$.

| Input | Scenario | Sel. coeff. $s^-$ Bias | Sel. coeff. $s^-$ RMSE | Sel. coeff. $s^+$ Bias | Sel. coeff. $s^+$ RMSE |
|---|---|---|---|---|---|
| GL | A | 0.00258 | 0.00438 | 0.00358 | 0.00646 |
| | B | 0.00111 | 0.00411 | 0.00100 | 0.00507 |
| | C | 0.00131 | 0.00387 | 0.00167 | 0.00627 |
| | D | 0.00067 | 0.00369 | −0.00018 | 0.00592 |
| | E | 0.00042 | 0.00372 | −0.00005 | 0.00541 |
| | F | 0.00056 | 0.00358 | −0.00051 | 0.00577 |
| CG | A | 0.00279 | 0.00467 | 0.00502 | 0.00790 |
| | B | 0.00159 | 0.00439 | 0.00333 | 0.00646 |
| | C | 0.00148 | 0.00403 | 0.00412 | 0.00733 |
| | D | 0.00080 | 0.00399 | 0.00224 | 0.00624 |
| | E | 0.00051 | 0.00368 | 0.00144 | 0.00548 |
| | F | 0.00053 | 0.00359 | 0.00058 | 0.00594 |

(b) Scenario 2: $s^- < 0$ and $s^+ = s^-$.

Table S9: Mean bias and RMSE in MAP estimates of the selection coefficients across different input types, data qualities and selection scenarios, corresponding to Figure S9. Mean bias and RMSE are calculated with 200 replicates for each combination. Data qualities (scenarios A–F) are described in Table 1, and selection scenarios (scenarios 1–9) are described in Table 2. GL and CG are shorthands separately for genotype likelihood and called genotype.

| Input | Scenario | Sel. coeff. $s^-$ Bias | Sel. coeff. $s^-$ RMSE | Sel. coeff. $s^+$ Bias | Sel. coeff. $s^+$ RMSE |
|---|---|---|---|---|---|
| GL | A | 0.00503 | 0.00777 | −0.00537 | 0.00896 |
| | B | 0.00214 | 0.00585 | −0.00263 | 0.00638 |
| | C | 0.00244 | 0.00547 | −0.00331 | 0.00725 |
| | D | 0.00128 | 0.00499 | −0.00165 | 0.00593 |
| | E | 0.00038 | 0.00462 | −0.00104 | 0.00558 |
| | F | 0.00052 | 0.00432 | −0.00018 | 0.00563 |
| CG | A | 0.00690 | 0.00981 | −0.00737 | 0.01115 |
| | B | 0.00455 | 0.00846 | −0.00533 | 0.00914 |
| | C | 0.00409 | 0.00702 | −0.00525 | 0.00928 |
| | D | 0.00270 | 0.00614 | −0.00291 | 0.00695 |
| | E | 0.00130 | 0.00503 | −0.00173 | 0.00586 |
| | F | 0.00106 | 0.00432 | −0.00169 | 0.01681 |

(c) Scenario 3: $s^- < 0$ and $s^+ > s^-$.

| Input | Scenario | Sel. coeff. $s^-$ Bias | Sel. coeff. $s^-$ RMSE | Sel. coeff. $s^+$ Bias | Sel. coeff. $s^+$ RMSE |
|---|---|---|---|---|---|
| GL | A | −0.00023 | 0.00347 | 0.00551 | 0.00834 |
| | B | 0.00028 | 0.00348 | 0.00191 | 0.00528 |
| | C | 0.00022 | 0.00366 | 0.00265 | 0.00571 |
| | D | 0.00064 | 0.00362 | −0.00027 | 0.00508 |
| | E | −0.00001 | 0.00338 | 0.00079 | 0.00432 |
| | F | 0.00063 | 0.00368 | −0.00035 | 0.00424 |
| CG | A | −0.00058 | 0.00366 | 0.00772 | 0.01087 |
| | B | −0.00013 | 0.00381 | 0.00483 | 0.00753 |
| | C | −0.00012 | 0.00360 | 0.00496 | 0.00793 |
| | D | 0.00023 | 0.00374 | 0.00243 | 0.00598 |
| | E | −0.00006 | 0.00336 | 0.00183 | 0.00546 |
| | F | 0.00057 | 0.00371 | 0.00022 | 0.00412 |

(d) Scenario 4: $s^- = 0$ and $s^+ < s^-$.

Table S9: Mean bias and RMSE in MAP estimates of the selection coefficients across different input types, data qualities and selection scenarios, corresponding to Figure S9, continued.

| Input | Scenario | Sel. coeff. $s^-$ | | Sel. coeff. $s^+$ | |
|---|---|---|---|---|---|
| | | Bias | RMSE | Bias | RMSE |
| GL | A | 0.00020 | 0.00389 | 0.00026 | 0.00300 |
| | B | −0.00001 | 0.00437 | 0.00044 | 0.00337 |
| | C | 0.00015 | 0.00368 | 0.00022 | 0.00292 |
| | D | 0.00015 | 0.00422 | 0.00040 | 0.00330 |
| | E | −0.00003 | 0.00391 | 0.00028 | 0.00301 |
| | F | −0.00007 | 0.00380 | 0.00040 | 0.00328 |
| CG | A | 0.00014 | 0.00370 | 0.00023 | 0.00285 |
| | B | −0.00011 | 0.00425 | 0.00036 | 0.00315 |
| | C | 0.00016 | 0.00348 | 0.00017 | 0.00277 |
| | D | 0.00017 | 0.00421 | 0.00033 | 0.00326 |
| | E | 0.00002 | 0.00380 | 0.00025 | 0.00293 |
| | F | −0.00022 | 0.00375 | 0.00062 | 0.00314 |

(e) Scenario 5: $s^- = 0$ and $s^+ = s^-$.

| Input | Scenario | Sel. coeff. $s^-$ | | Sel. coeff. $s^+$ | |
|---|---|---|---|---|---|
| | | Bias | RMSE | Bias | RMSE |
| GL | A | −0.00031 | 0.00368 | −0.00460 | 0.00722 |
| | B | −0.00084 | 0.00409 | −0.00134 | 0.00548 |
| | C | −0.00054 | 0.00394 | −0.00195 | 0.00571 |
| | D | −0.00061 | 0.00411 | 0.00021 | 0.00548 |
| | E | −0.00060 | 0.00365 | −0.00034 | 0.00494 |
| | F | −0.00086 | 0.00397 | 0.00037 | 0.00427 |
| CG | A | 0.00000 | 0.00354 | −0.00656 | 0.00952 |
| | B | −0.00033 | 0.00420 | −0.00453 | 0.00756 |
| | C | −0.00032 | 0.00385 | −0.00422 | 0.00749 |
| | D | −0.00029 | 0.00408 | −0.00160 | 0.00523 |
| | E | −0.00039 | 0.00371 | −0.00142 | 0.00497 |
| | F | −0.00071 | 0.00390 | −0.00001 | 0.00422 |

(f) Scenario 6: $s^- = 0$ and $s^+ > s^-$.

Table S9: Mean bias and RMSE in MAP estimates of the selection coefficients across different input types, data qualities and selection scenarios, corresponding to Figure S9, continued.

| Input | Scenario | Sel. coeff. $s^-$ Bias | Sel. coeff. $s^-$ RMSE | Sel. coeff. $s^+$ Bias | Sel. coeff. $s^+$ RMSE |
|---|---|---|---|---|---|
| GL | A | −0.00558 | 0.00834 | 0.00585 | 0.00951 |
| | B | −0.00259 | 0.00619 | 0.00200 | 0.00611 |
| | C | −0.00265 | 0.00580 | 0.00195 | 0.00624 |
| | D | −0.00128 | 0.00529 | 0.00021 | 0.00593 |
| | E | −0.00064 | 0.00452 | 0.00002 | 0.00569 |
| | F | −0.00067 | 0.00475 | 0.00034 | 0.00552 |
| CG | A | −0.00774 | 0.01066 | 0.00825 | 0.01236 |
| | B | −0.00506 | 0.00837 | 0.00507 | 0.00953 |
| | C | −0.00432 | 0.00741 | 0.00406 | 0.00805 |
| | D | −0.00304 | 0.00670 | 0.00272 | 0.00693 |
| | E | −0.00158 | 0.00522 | 0.00083 | 0.00564 |
| | F | −0.00111 | 0.00480 | 0.00078 | 0.00561 |

(g) Scenario 7: $s^- > 0$ and $s^+ < s^-$.

| Input | Scenario | Sel. coeff. $s^-$ Bias | Sel. coeff. $s^-$ RMSE | Sel. coeff. $s^+$ Bias | Sel. coeff. $s^+$ RMSE |
|---|---|---|---|---|---|
| GL | A | −0.00295 | 0.00521 | −0.00383 | 0.00721 |
| | B | −0.00131 | 0.00415 | −0.00093 | 0.00596 |
| | C | −0.00104 | 0.00371 | −0.00219 | 0.00628 |
| | D | −0.00066 | 0.00389 | 0.00001 | 0.00547 |
| | E | −0.00090 | 0.00385 | −0.00033 | 0.00598 |
| | F | −0.00092 | 0.00359 | 0.00079 | 0.00572 |
| CG | A | −0.00342 | 0.00562 | −0.00537 | 0.00862 |
| | B | −0.00164 | 0.00476 | −0.00325 | 0.00731 |
| | C | −0.00161 | 0.00409 | −0.00407 | 0.00740 |
| | D | −0.00093 | 0.00397 | −0.00170 | 0.00602 |
| | E | −0.00100 | 0.00380 | −0.00113 | 0.00633 |
| | F | −0.00105 | 0.00372 | 0.00014 | 0.00602 |

(h) Scenario 8: $s^- > 0$ and $s^+ = s^-$.

Table S9: Mean bias and RMSE in MAP estimates of the selection coefficients across different input types, data qualities and selection scenarios, corresponding to Figure S9, continued.

| Input | Scenario | Sel. coeff. $s^-$ Bias | Sel. coeff. $s^-$ RMSE | Sel. coeff. $s^+$ Bias | Sel. coeff. $s^+$ RMSE |
|---|---|---|---|---|---|
| GL | A | −0.00167 | 0.00412 | −0.00713 | 0.00982 |
| | B | −0.00124 | 0.00434 | −0.00294 | 0.00792 |
| | C | −0.00142 | 0.00380 | −0.00284 | 0.00744 |
| | D | −0.00019 | 0.00350 | −0.00006 | 0.00687 |
| | E | −0.00047 | 0.00332 | −0.00082 | 0.00597 |
| | F | −0.00038 | 0.00365 | −0.00068 | 0.00612 |
| CG | A | −0.00183 | 0.00450 | −0.00984 | 0.01226 |
| | B | −0.00128 | 0.00454 | −0.00817 | 0.01194 |
| | C | −0.00136 | 0.00389 | −0.00671 | 0.00972 |
| | D | 0.00017 | 0.00358 | −0.00397 | 0.00885 |
| | E | −0.00034 | 0.00349 | −0.00268 | 0.00615 |
| | F | −0.00034 | 0.00360 | −0.00171 | 0.00719 |

(i) Scenario 9: $s^- > 0$ and $s^+ > s^-$.

Table S9: Mean bias and RMSE in MAP estimates of the selection coefficients across different input types, data qualities and selection scenarios, corresponding to Figure S9, continued.

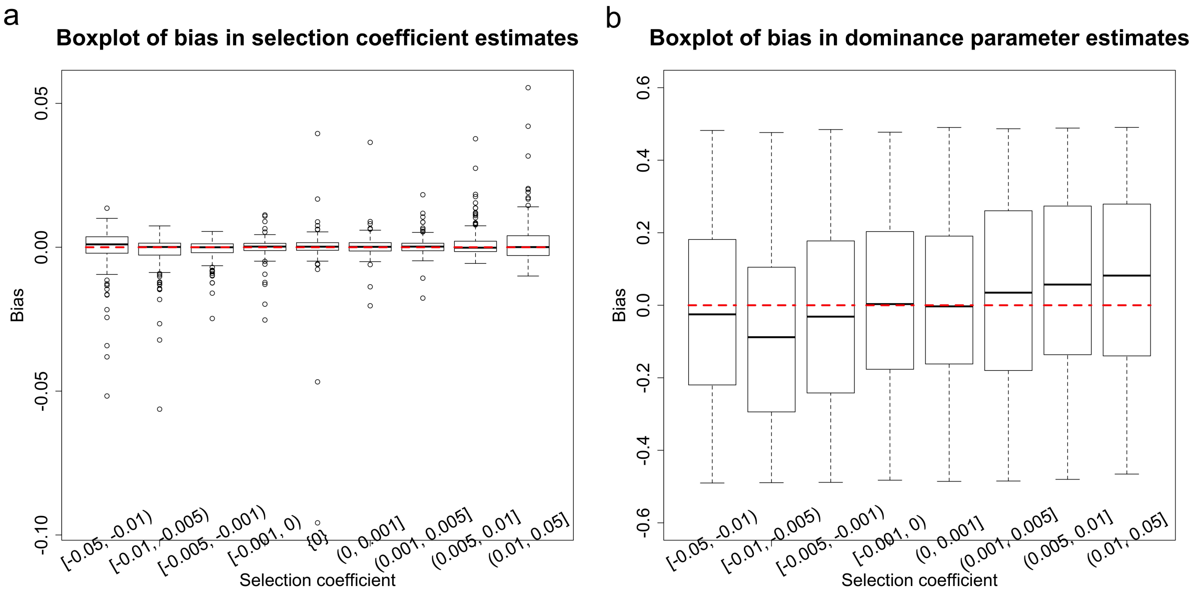


Figure S11: Empirical distributions for the bias in MAP estimates of the selection coefficient and dominance parameter across different ranges of the selection coefficient $s$ with the parameters $\phi = 0.85$ and $\psi = 1$ (*i.e.*, scenario D in Table 1. The simulated datasets for each range of the selection coefficient are the same as those in Figure 4. Boxplots for the bias in estimation of (a) the selection coefficient and (b) the dominance parameter.

| Sel. coeff. $s$ | Sel. coeff. $s$ | | Dom. param. $h$ | |
|---|---|---|---|---|
| | Bias | RMSE | Bias | RMSE |
| $s \in [-0.050, -0.010)$ | $-0.00035$ | 0.00734 | $-0.01851$ | 0.25650 |
| $s \in [-0.010, -0.005)$ | $-0.00163$ | 0.00658 | $-0.08433$ | 0.26988 |
| $s \in [-0.005, -0.001)$ | $-0.00087$ | 0.00367 | $-0.03049$ | 0.27109 |
| $s \in [-0.001, 0)$ | $-0.00016$ | 0.00351 | 0.00004 | 0.25984 |
| $s \in \{0\}$ | $-0.00002$ | 0.00845 | N/A | N/A |
| $s \in (0, 0.001]$ | 0.00024 | 0.00383 | 0.00905 | 0.24026 |
| $s \in (0.001, 0.005]$ | 0.00004 | 0.00323 | 0.03563 | 0.27933 |
| $s \in (0.005, 0.010]$ | 0.00125 | 0.00531 | 0.06200 | 0.26997 |
| $s \in (0.010, 0.050]$ | 0.00165 | 0.00779 | 0.07517 | 0.26429 |

Table S10: Mean bias and RMSE in MAP estimates of the selection coefficient and dominance parameter across different ranges of the selection coefficient $s$ with the parameters $\phi = 0.85$ and $\psi = 1$ (*i.e.*, scenario D in Table 1), corresponding to Figure S11. Mean bias and RMSE are calculated with 200 replicates for each case.

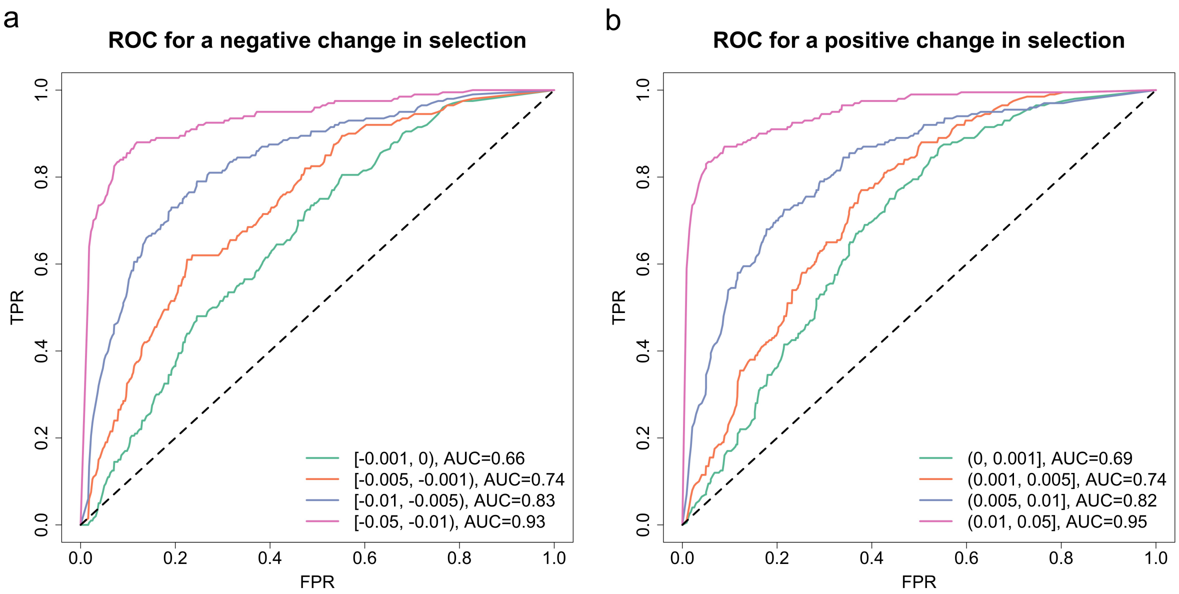


Figure S12: ROC curves for testing a change in selection across different ranges of the selection change $\Delta s$ with the parameters $\phi = 0.85$ and $\psi = 1$ (*i.e.*, scenario D in Table 1. The AUC value for each curve is summarised. The simulated datasets for each range of the selection change are the same as those in Figure 6. ROC curves for (a) a negative change in selection and (b) a positive change in selection. Compared to Figure 6, the dominance parameter $h$ is estimated rather than prespecified.

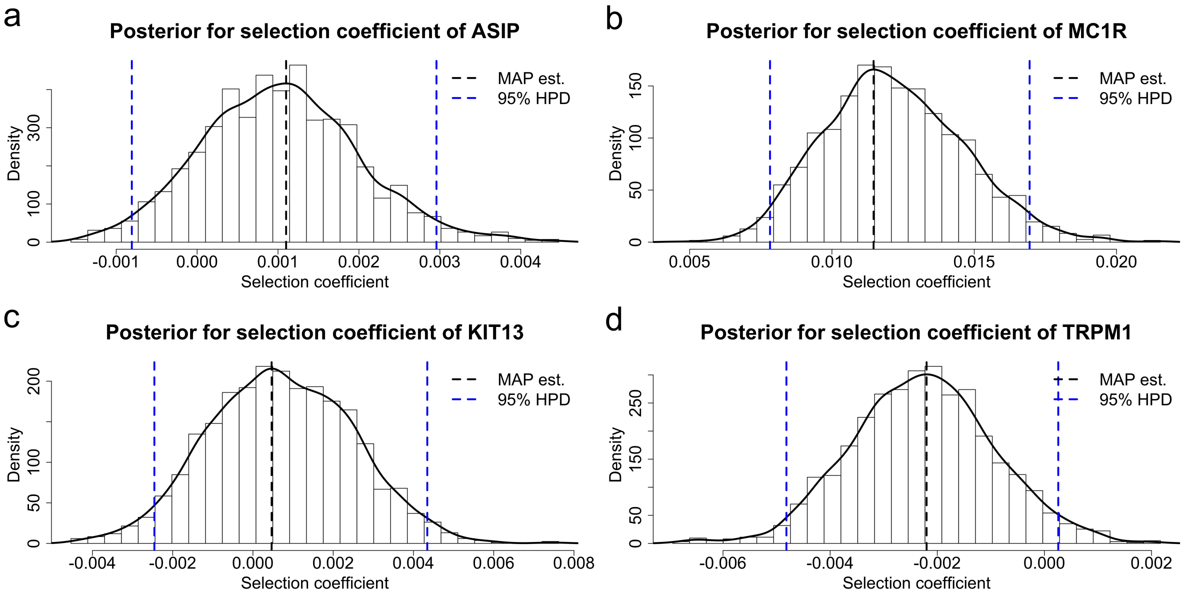


Figure S13: Posteriors for the selection coefficients of (a) *ASIP*, (b) *MC1R*, (c) *KIT13* and (d) *TRPM1*. We assume that the selection coefficient of each gene is fixed over time, and run our procedure on the ancient horse samples presented in Table S4 with the same sample exclusion criteria for each gene.

| Gene | Sel. coeff. | MAP est. | 95% HPD | Prob. for -ve. | Prob. for +ve. |
|---|---|---|---|---|---|
| *ASIP* | $s$ | 0.00110 | $[-0.00081, 0.00296]$ | 0.1375 | 0.8625 |
| *MC1R* | $s$ | 0.01146 | $[\ \ 0.00782, 0.01695]$ | 0.0000 | 1.0000 |
| *KIT13* | $s$ | 0.00047 | $[-0.00246, 0.00434]$ | 0.3540 | 0.6460 |
| *TRPM1* | $s$ | $-0.00220$ | $[-0.00482, 0.00026]$ | 0.9565 | 0.0435 |

Table S11: MAP estimates of the selection coefficients with their 95% HPD intervals, as well as posterior probabilities for negative selection and positive selection, for *ASIP*, *MC1R*, *KIT13* and *TRPM1*, corresponding to Figure S13. Prob. for -ve. and +ve. stand for posterior probabilities for negative selection and positive selection, respectively.
